# Non-invasive detection of tissue clonal mosaicism in chronic disease

**DOI:** 10.64898/2026.11.20.767624

**Authors:** Srinivas Rajagopalan, Boyuan Li, Alexandre Pellan Cheng, Maria Torra I Benach, Natalia Brzozowska, John Zinno, Andrew R. D’Avino, Theo Botella, William Driscoll, Anders Rasmussen, Wesley S. Lu, Joe Pelt, Rob Furatero, Alexi Runnels, Erica Wolin, Itai Rusinek, Ariel Jaimovich, Cristina Cotobal Martin, Wendy Garri, Miles Parkes, Emily Ashwin, James Robineau, Min Zhu, Ashwini Arvind, Andrew Shubin, Mengmeng Wang, David Hsieh, Amit Singal, Madhukar Patel, Catherine Potenski, Adam J. Widman, Foad J. Rouhani, Matthew Hoare, Maria T. Abreu, Peter J. Campbell, Tim Raine, Carl A. Anderson, Eleanor Barnes, Hao Zhu, Dan A. Landau

**Affiliations:** New York Genome Center, New York, NY, USA; Department of Systems and Computational Biomedicine, Weill Cornell Medicine, New York, NY, USA; Physiology, Biophysics and Systems Biology Graduate Program, Weill Cornell Medicine, New York, NY, USA; Children’s Research Institute, Departments of Pediatrics and Internal Medicine, Simmons Comprehensive Cancer Center, Center for Regenerative Science and Medicine, University of Texas Southwestern Medical Center, Dallas, TX, USA; Département de génie des systèmes, École de technologie supérieure, Montréal, QC, Canada; Axe Cancer, Centre de Recherche du Centre Hospitalier de l’Université de Montréal (CRCHUM), Montréal, QC, Canada; Wellcome Sanger Institute, Hinxton, UK; Wellcome-MRC Cambridge Stem Cell Institute, Cambridge Biomedical Campus, Cambridge, UK; Ultima Genomics, Fremont, California, USA; Cambridge University Hospitals, Cambridge, UK; Nuffield Department of Medicine, University of Oxford, Oxford, UK; Department of Internal Medicine, Division of Digestive and Liver Diseases, University of Texas Southwestern Medical Center, Dallas, TX, USA; Division of Surgical Transplantation, Department of Surgery, University of Texas Southwestern Medical Center, Dallas, TX, USA; LifeShare Network Inc., Oklahoma City, OK, USA; Department of Internal Medicine, Division of Hematology-Oncology, Simmons Comprehensive Cancer Center, University of Texas Southwestern Medical Center, Dallas, TX, USA; Memorial Sloan Kettering Cancer Center, New York, NY, USA; Tissue Regeneration and Clonal Evolution Laboratory, The Francis Crick Institute, London, UK; Roger Williams Institute of Liver Studies, School of Immunology &Microbial Sciences, Faculty of Life Sciences and Medicine, King’s College London, Foundation for Liver Research and King’s College Hospital, London, UK; F. Widjaja Inflammatory Bowel Disease Institute, Cedars-Sinai Medical Center, Los Angeles, CA, USA

## Abstract

Liquid biopsy has transformed our ability to non-invasively interrogate the somatic genome, but its reach has been largely confined to fetal genetics in prenatal testing and tumor genetics in cancer. Clonal mosaicism (CM) pervades non-malignant solid tissues, where somatic driver mutations expand under specific selective pressures and may shape the course of chronic illness. Access to this biology has been limited to invasive tissue biopsies, restricting the field to small cohorts and single time points, and has precluded large-scale, clinically-linked analyses. Here, ultra-sensitive cell-free DNA (cfDNA) sequencing of patients with chronic liver disease revealed genome-wide somatic variants from matched liver biopsies at plasma fractions comparable to those of cancers (10^−4^–10^−5^), with signal corresponding to variants from spatially and phylogenetically related clones. A biopsy-naive targeted panel detected pathogenic variants in chronic liver disease driver genes in 81.5% of patients with chronic liver disease, undetectable in controls, with stage-specific enrichment of ACVR2A in early disease and clearance of liver-derived signal following transplant. Extending the framework to inflammatory bowel disease (IBD), we detect clonal expansions of IBD driver genes in active ulcerative colitis, and observe greater detection in patients with pancolitis versus more localized disease. Together, these results open a new domain for cfDNA genetics beyond cancer and prenatal genetics, the non-invasive interrogation of somatic evolution in chronic disease, and establish a framework for the large-cohort, longitudinal, clinically linked studies of solid-tissue CM that has been inaccessible to the field.

## Introduction

Over the past two decades, the analysis of cell-free DNA circulating in plasma has transformed non-invasive access to the somatic genome. Two clinical domains have driven this transformation. In prenatal genetics, the detection of fetal cell-free DNA in maternal circulation enabled non-invasive prenatal testing (NIPT) at population scale^1,2^. In oncology, circulating tumor DNA (ctDNA) has become a central tool for cancer detection, residual disease monitoring, and treatment response^3–5^. In both domains, cell-free DNA succeeded via detection of a genetically distinct cell population shedding DNA into plasma.

A third, far broader substrate of human somatic variation has emerged in recent years: clonal mosaicism of non-malignant tissues. Somatic mutations accumulate throughout life in every tissue, and a subset confers clonal fitness advantages that drive expansions visible across normal and diseased human tissues, including blood^6,7^, esophagus^8,9^, colon^10^, endometrium^11,12^, bladder^13^, skin^14^, and liver^15–18^. This clonal mosaicism (CM) is increasingly recognized as a fundamental feature of human tissue biology^19^.

In chronic disease, CM is not incidental. Recurrent positively-selected clones expand specifically in response to disease-associated environments: loss-of-function variants in lipid metabolism genes such as *CIDEB* and *GPAM* expand in the steatotic liver of metabolic dysfunction-associated steatotic liver disease (MASLD)^15,16^. The biological significance of these drivers is further supported by germline genetics: rare loss-of-function and missense variants in *CIDEB* carried by ∼0.7% of the population are associated with reduced aminotransferase levels and 33% lower odds of liver disease of any cause, including cirrhosis, with a corresponding reduction in NAFLD activity score (NAS) and protection against lipid droplet accumulation in hepatoma cell lines^20^. Immune-modulatory variants in genes such as *NFKBIZ, PIGR*, and *TRAF3IP2* expand in the inflamed colonic epithelium of ulcerative colitis^21–23^. These clones plausibly shape disease trajectory, either by mitigating pathological pressures (loss of *CIDEB* reducing lipid droplet formation in MASLD) or by dampening inflammatory signaling (loss of *IL17RA* in IBD), raising the possibility that the specific clonal composition of a diseased tissue is a biomarker, and potentially a modifier, of chronic disease.

The clinical importance of somatic clonal expansions was established most clearly in blood. The recognition that clonal hematopoiesis of indeterminate potential (CHIP) accumulates with age and associates with cardiovascular, metabolic, inflammatory, and malignant outcomes transformed an obscure observation into a major axis of translational research, ultimately influencing how we think about aging, risk stratification, and cancer predisposition^6,7,24,25^. Crucially, blood can be sampled longitudinally, and at scale, allowing large-cohort epidemiology and clinical linkage. In contrast, solid-tissue CM studies have been constrained to tens of patients, often dependent on surgical explants, rapid autopsies, or esearch-only biopsies^16,26^. Compounding the problem, solid tissues are spatially organized: unlike blood, a single biopsy samples only a small patch of a larger clonal landscape, so even when tissue is available, it captures a fragment of the mosaic.

Recent work has begun to show that cell-free DNA carries information beyond cancer and prenatal genetics: methylation-based deconvolution detects non-malignant tissue damage from β-cell death in diabetes to hepatocyte injury^27,28^, and ultra-sensitive sequencing has begun to recover cell-free DNA from premalignant lesions^29,30^. Motivated by these signals, we hypothesized that the somatic clonal expansions of chronically diseased solid tissues could be detected directly in plasma at clone resolution. Here we establish a liquid biopsy framework for the non-invasive detection of solid-tissue clonal mosaicism in chronic disease. Using cell-free DNA from patients with liver diseases, including MASLD, and inflammatory bowel disease (IBD), two mechanistically distinct chronic illnesses with well-characterized driver landscapes, we demonstrate that disease-associated somatic clones shed detectable signal into plasma; that biopsy-naive targeted sequencing can detect this signal across large cohorts without matched tissue; that the signal tracks disease stage, responds to curative intervention, and resolves multi-organ clonal activity from a single plasma draw. We thus present a framework for the study of solid-tissue CM as a clinical feature of chronic disease. We envision a longitudinal, clinically-linked, population-scale exploration of CM in chronic disease and a new horizon for precision medicine based on somatic genetics.

## Results

### Whole-genome sequencing (WGS) of matched tissue and plasma cell-free DNA establishes the shedding of cell-free DNA from diseased liver clonal mosaicism (CM)

To explore the hypothesis that solid tissue CM can be detected non-invasively in the plasma, we first set out to develop highly sensitive approaches, focusing on CM in liver disease^15^. We began with a deeply characterized MASLD cohort: three patients (LD-001, LD-002, LD-003) for whom matched cell-free DNA was available alongside dense whole-liver laser capture microdissection (LCM) sampling, with 220-284 microdissections drawn from eight anatomical segments per liver^15^ (**Fig. 1A, Supplementary Table 1, Supplementary Table 2**). Notably, cell-free DNA from these three patients was isolated from serum rather than plasma, because only serum samples were available. Because serum-derived cell-free DNA is enriched for higher-molecular-weight DNA fragments and may exhibit reduced tissue contributions relative to plasma-derived cell-free DNA^31^, standard SPRI bead cleanup was used to retain short, mononucleosomal fragments (Methods).

**Figure 1:**
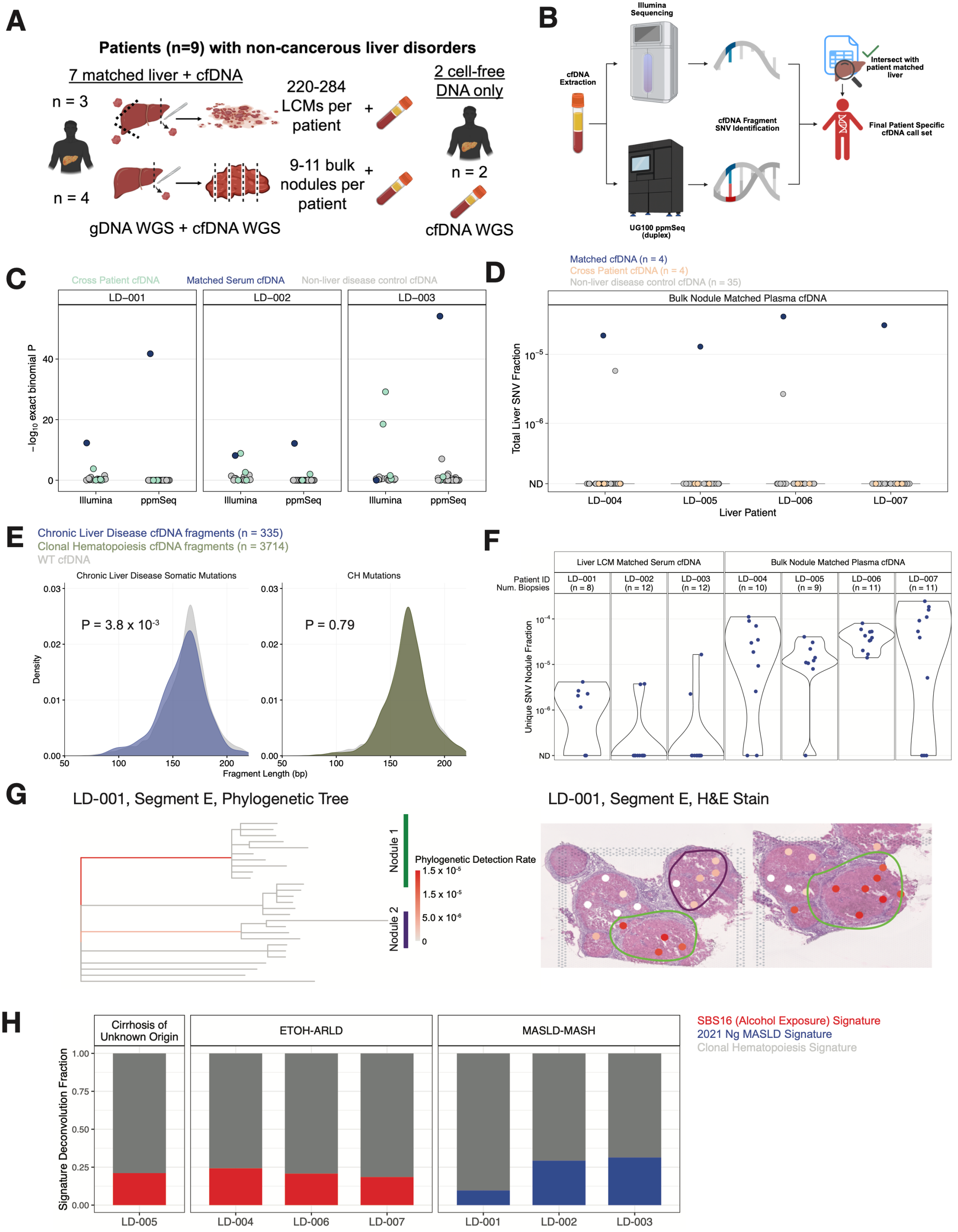
Non-cancerous liver disease can be detected non-invasively through plasma cell-free DNA sequencing. **A**, Schematic of liver disease cohort comprising n = 7 patients with matched liver biopsy and cell-free DNA samples and n = 2 patients with only cell-free DNA samples. Liver disease patients include those with Metabolic Dysfunction-Associated Steatotic Liver Disease (MASLD); alcohol-related cirrhosis, Metabolic Dysfunction-Associated Steatohepatitis (MASH) or cirrhosis of unknown etiology (cryptogenic cirrhosis; Supplementary Table 1). LCM, laser capture microdissection. WGS, whole-genome sequencing. gDNA, genomic DNA. **B**, Workflow outlining the sequencing and analysis strategy to obtain patient-specific profiles to non-invasively detect liver-derived somatic variants from cell-free DNA (cfDNA). **C**, One-sided binomial test P values for detected patient-specific SNVs identified from matched laser capture microdissection liver biopsies in cell-free DNA whole-genome sequences (matched serum cell-free DNA) compared to cross-patient SNV profiles in cell-free DNA whole-genome sequences (n = 4 serum samples for Illumina; n = 2 serum samples for ppmSeq) and cell-free DNA from non-liver disease controls (n = 13 for Illumina; n = 35 for ppmSeq) in three patients with MASLD. Cell-free DNA was sequenced using either Illumina or ppmSeq. **D**, Fraction of patient-specific SNVs identified from matched large cirrhotic liver nodule biopsies detected in cell-free DNA whole-genome sequences (ppmSeq) across n = 4 liver disease patients, compared to cross-patient SNV profiles from cell-free DNA (n = 4 plasma samples) and cell-free DNA from non-liver disease controls (n = 35 plasma samples). Detection rate is the number of unique supporting mixed cell-free DNA reads divided by the product of the mean sequencing coverage of the sample, the number of single-nodule SNVs interrogated, and the mixed-read rate of the sample. A lasso was applied to all three sample classes, in which a sample with only one supporting read across the entire patient-specific variant set had its detection rate set to zero. **E**, Left, fragment length distribution analysis of cell-free DNA for fragments carrying liver-confirmed variants identified from matched liver biopsy (Chronic Liver Disease cfDNA fragments) compared to wild-type (WT) cell-free DNA without liver-confirmed fragments. Right, fragment length distribution analysis of cell-free DNA for fragments carrying clonal hematopoiesis (Clonal Hematopoiesis cfDNA fragments) variants compared to wild-type (WT) cell-free DNA without clonal hematopoiesis fragments. bp, basepairs. Analysis restricted to validation-cohort plasma cell-free DNA. Two-sided Wilcoxon test. **F**, Fraction of unique biopsy-specific SNVs identified from matched laser capture microdissection liver biopsies (found in spatial proximity from the same biopsy) or large cirrhotic liver nodule biopsies detected in cell-free DNA from either plasma or serum using whole-genome sequences (ppmSeq) across n = 7 chronic liver disease patients. A lasso was applied, in which a sample with only one supporting read across the entire biopsy-specific variant set had its detection rate set to zero. Number of biopsies per patient is indicated. **G**, Phylogenetic tree (left) constructed using somatic SNVs^15^ across nodules from liver Segment E from patient LD-001, with detection rate of microdissection-specific SNVs in cell-free DNA mapped onto a hematoxylin and eosin (H&E)-stained light micrograph of liver Segment E (right, two z stack images of the same liver segment). Two nodules with different levels of SNV detection in the cell-free DNA are marked in green and purple on the phylogeny branches and micrograph. **H**, Trinucleotide single-base substitution signature analysis of cell-free DNA from cirrhosis of unknown etiology (n = 1 patient) sample, alcohol-related liver disease (ETOH-ARLD; n = 3 patients) samples and MASLD-MASH (n = 3 patients) samples. Mutation signatures were extracted from dual strand whole-genome sequencing (ppmSeq) of plasma cell-free DNA. Plasma signatures were fit to a custom catalog comprising the following signatures: clonal hematopoiesis^47^, SBS16 (alcohol exposure) and a *de novo* MASLD signature from Ng et al.^15^. Fractions of SBS16, MASLD signature, and clonal hematopoiesis signatures are displayed for each patient.

We adopted a tumor-informed WGS framework, in which a per-patient set of somatic variants is defined from WGS of the tissue of interest and the matched plasma WGS is then mined specifically for cell-free DNA fragments supporting these pre-defined variants, an approach previously shown to enable the detection of circulating tumor DNA shedding even from precancerous adenomas and pT1 colorectal carcinomas^29^. Here, we applied the same logic in the context of chronic liver disease. LCM samples harbored a median of 1,382 (range 97 to 8,482) somatic single-nucleotide variants (SNVs), fewer than the ∼9,000 somatic mutations typical of hepatocellular carcinoma whole genomes^32^. Pooled across the liver, this gave per-patient sets of 144,201 (LD-001), 361,601 (LD-002), and 808,948 (LD-003) SNVs against which matched cell-free DNA was queried^15^; the union is smaller than the per-LCM sum as some SNVs are shared across proximal LCMs. We then performed deep Illumina WGS (∼300x) on the matched cell-free DNA from these three patients, with read-level quality filters applied to suppress sequencing artifacts^29^ (Methods). For each patient, we queried the cell-free DNA for fragments carrying SNVs from the matched patient-specific liver variant call format (VCF) file. We compared these matched detection rates to those obtained from unmatched chronic liver disease patient cell-free DNA and cell-free DNA from non-liver disease controls^29,33^ (**Fig. 1B**). To address the potential confounder of clonal hematopoiesis-associated mutations that may pervade the liver through immune cell infiltration^34^, we removed SNVs present across multiple liver segments or biopsies from each patient-specific VCF (151 in LD-001, 353 in LD-002, 399 in LD-003).

Recurrence across spatially distinct samples is unlikely for identical liver-disease-specific variants (outside of known hotspot variants such as *FOXO1p*.*S22W*) and is a known signature of clonal hematopoiesis^17^. Consistent with this assignment, the excluded mutations had markedly elevated cell-free DNA variant allele frequencies (VAFs) relative to retained liver-specific variants. We generated binomial test P values for coverage normalized detection rates of patient-specific liver-derived somatic SNVs in cell-free DNA (Methods). The 300X Illumina data showed statistically significant increase in detection scores for patient-matched comparisons in two of the three patients (LD-001 and LD-002), but with limited separation from the cross-patient and non-liver disease control distributions, while the third patient (LD-003) showed no separation from background (**Fig. 1C, Supplementary Fig. 1**). The narrow margin above background in the two positive patients, combined with the absent signal in the third, was suggestive of liver-derived shedding into plasma but did not allow us to confidently distinguish true signal from library and degraded-DNA artifacts inherent to cell-free DNA sequencing.

When the two strands of highly degraded cell-free DNA are independently sequenced, lesions from library prep and DNA degradation are read as variants^35^. Paired plus-minus strand sequencing (ppmSeq), a recently developed technology based on Ultima Genomics (UG) sequencing, provides cost-efficient double-strand DNA sequencing, suppressing single-strand library artifacts and enabling highly sensitive variant detection at low error rates at genome scale^36^. We therefore reanalyzed the cell-free DNA from the three MASLD patients (LD-001, LD-002, LD-003) using ppmSeq, with dual-strand fractions of 0.30 - 0.42 (**Supplementary Table 3**). ppmSeq cell-free DNA libraries were also generated for a separate set of non-liver disease controls to serve as the comparator for these analyses (dual-strand fractions of 0.405 - 0.470; **Supplementary Table 3**). As with the Illumina analysis above, we queried the ppmSeq cell-free DNA for fragments carrying SNVs from the matched per-patient liver SNV set, retaining only variants supported by both strands of the original molecule (termed mixed reads), and compared detection rates against unmatched chronic liver cell-free DNA and non-liver disease control cell-free DNA^29^ (Methods). The ppmSeq data on these same three patients showed substantially higher detection scores in patient-matched samples than in unmatched cross-patient or non-liver disease control samples (**Fig. 1C**). These data, using molecular error correction based on dual strand (duplex) sequencing, confirmed that liver CM mutations can be confidently detected in the circulation.

To validate these findings, we collected additional matched liver explants and plasma samples from four cirrhosis patients (LD-004, LD-005, LD-006, LD-007; three with alcohol-related etiology, one cryptogenic cirrhosis; **Fig. 1A, Supplementary Table 1**). Unlike the initial three-patient cohort, where cell-free DNA was isolated from serum, here we used plasma-derived cell-free DNA, which is less impacted by high-molecular-weight contamination^31^. Each explant was divided into discrete nodules (visible regions of abnormal growth) and subjected to UG ppmSeq to a target coverage of 100x (n = 9-11 nodules per patient; **Supplementary Table 3**). Somatic SNVs were called from the bulk-nodule WGS using both dual-strand (duplex) and non-dual strand reads, with machine-learning-based read-level denoising and a binomial test against an in-house UG whole-genome panel (n = 91 WGS from normal tissue) for error polishing^35,37,38^. Germline and clonal hematopoiesis variants were then removed, and only variants present in a single biopsy were retained, yielding the per-patient variant set (Methods). The average number of mutations per liver nodule was 2,278 (range 147 to 18,097), consistent with the somatic SNV burden from previous reports^15^ (**Supplementary Fig. 2**). We performed Ultima ppmSeq on the matched plasma cell-free DNA, with a target coverage of 200x. Analogous to our previous analysis, we queried our per-patient variant sets against the matched plasma, the unmatched plasma, and non-liver disease controls sequenced with ppmSeq. Across these four patients, we detected strong signal in the patient-matched comparisons, with total liver SNV fractions as high as 10^−5^-10^−4^ (**Fig. 1D**), comparable to circulating tumor DNA fractions reported for cancer^29^.

In cancer, circulating tumor DNA is known to have a shorter fragment length than the general cell-free DNA pool^39–41^, providing a useful feature that can be leveraged to discriminate circulating tumor DNA from background cell-free DNA^29^. Recent work additionally demonstrated that cell-free DNA fragmentomics models can be used to develop liquid biopsies capable of identifying early chronic liver disease and fibrosis in cirrhosis using standard whole-genome sequencing^42^. Using the ppmSeq plasma data from the four cirrhosis patients, where fragment-length distributions are better preserved than in serum, we tested whether liver-derived fragments in our cohort exhibited a similar fragment-length shift. As a positive control, we sequenced pre-treatment plasma cell-free DNA from a high-tumor-burden bladder cancer patient with matched tumor WGS, identified circulating tumor DNA fragments using the tumor VCF, and recovered the same short-fragment shift relative to wild-type fragments at the same loci (**Supplementary Fig. 3**). Analogously, we then classified cell-free DNA fragments as either cirrhotic liver-derived (bearing a patient-matched SNV identified from liver sequencing) or wild type (fragments of DNA not containing any SNVs identified in matched livers at the same locus of patient-matched SNVs) and assessed fragment length. In the ppmSeq plasma datasets from the four liver cirrhosis patients (LD-004 through LD-007), liver SNV-bearing fragments were overall shorter than wild-type fragments (P = 3.8 × 10^−3^; **Fig. 1E**, left). This finding was corroborated in the Illumina serum cell-free DNA data from the three MASLD patients (LD-001, LD-002 and LD-003), where MASLD SNV-bearing fragments were similarly shorter than wild-type fragments (P = 4.58 × 10^−5^; **Supplementary Fig. 4**). This is in contrast with clonal hematopoiesis fragments^43,44^, defined as those bearing SNVs shared across all nodules from a given patient^17^, where we observed no shift in size (P = 0.79; **Fig. 1E**, right). These data, showing that liver-derived cell-free DNA is shorter, suggest that the diseased liver environment may affect cell-free DNA fragment length, possibly through altered epigenetics, nucleosome positioning or nuclease activity^39,45^.

We next assessed whether cell-free DNA CM contribution varied across different regions of the liver. For this analysis, we identified SNVs that were on LCMs grouped together from the same biopsy (in the three MASLD patients LD-001, LD-002 and LD-003) or unique to individual explant nodules (in the four cirrhosis patients LD-004 to LD-007). We observed a range of CM cell-free DNA contribution that spanned almost three orders of magnitude (10^−3^-10^−^6) (**Fig. 1F**), indicating that cell-free DNA shedding is not uniform across the liver. To illustrate this variability, we revisited the published phylogenetic tree of clonal outgrowth from Ng et al.^15^, which groups SNVs across LCMs and mapped the cell-free DNA detection rate of each SNV group onto the spatial location of its LCMs. We focused on Segment E from patient LD-001, which was best powered for this analysis with the largest number of shedding LCMs on a single biopsy. SNVs originating from Nodule 1 (green circle, **Fig. 1G**) had strikingly higher detection rates in the cell-free DNA than SNVs specific to Nodule 2 (purple circle, **Fig. 1G**). We examined whether known drivers of chronic liver disease or differential mutational signatures could explain the difference in shedding between the two nodules. Neither nodule harbored chronic liver disease driver mutations, nor did either nodule carry a unique COSMIC signature absent from the other. The basis for region-specific shedding therefore remains an open question for future studies.

Liver biopsies sequenced with WGS represent only around 0.0001% of the 100 billion hepatocytes present in the typical liver. Many additional clones go unsampled by these biopsies but can still shed into the cell-free DNA pool. As our WGS surveys the entire genome, it may capture variants from these unsampled clones. Notably, dual strand sequencing with ppmSeq provides cell-free DNA mutation calls that are sufficiently error-free to read out mutational processes directly from plasma, as recently demonstrated for APOBEC mutagenesis and UV exposure in cancer^36,46^. Building on this, we interrogated our data for the detection of liver-related mutational signatures, without using the matched tissue sequencing. Two such processes are well-established in liver disease: SBS16, associated with alcohol exposure and observed in liver biopsies from patients with a history of alcohol use^15,16^, and a de novo MASLD signature previously identified and exclusively observed in the livers of MASLD patients^15^. We extracted mutational signatures from the cell-free DNA ppmSeq WGS of the three MASLD patients (LD-001 through LD-003) and the four cirrhosis patients (LD-004 through LD-007; three with alcohol-related cirrhosis and one cryptogenic), and fit these to a catalog comprising the clonal hematopoiesis signature (control)^47^, SBS16 (alcohol exposure), and the de novo MASLD signature^15^ (Methods). The de novo MASLD signature was observed only in the cell-free DNA of the three MASLD patients, and the SBS16 signature was observed in the cell-free DNA of alcohol-related liver disease (ETOH-ARLD) patients, as well as in the cryptogenic cirrhosis patient (**Fig. 1H**), demonstrating that we can detect disease exposure and signal in the cell-free DNA of liver disease patients even in the absence of patient-specific profiles obtained from matched liver sequencing, and further suggesting that cell-free DNA mutational signatures may in some cases reveal exposure history not captured by clinical etiology assignment.

These data establish that DNA fragments bearing somatic mutations from chronically diseased livers are shed into the circulation at detectable levels, with properties, including shorter fragment lengths and disease-specific mutational signatures, mirroring those of circulating tumor DNA in cancer, supporting the feasibility of non-invasive monitoring of chronic liver disease-derived variants from cell-free DNA.

### Deep targeted cell-free DNA sequencing enables detection of liver clonal mosaicism driver-gene mutations

Having established that liver-derived somatic variants can be detected in plasma cell-free DNA through WGS informed by matched liver biopsies, we next sought to develop an approach that could be applied to large clinical cohorts without requiring matched tissue, translating these findings into a scalable, non-invasive framework for larger cohorts and broader disease contexts by establishing a targeted sequencing approach. Recurrent somatic driver mutations in chronic liver disease have been identified through DNA sequencing of liver biopsies, primarily impacting lipid metabolism genes such as *FOXO1, CIDEB*, and *GPAM*^15^. The clonal architecture of the diseased liver, with many small clones scattered across the organ rather than a single dominant clone, is compatible with deep targeted sequencing of recurrently selected driver genes, which would capture variants from different clones across the liver. Detection of these variants in plasma cell-free DNA would enable scalable, non-invasive monitoring of chronic liver disease-associated clonal mosaicism and could be tested for clinical correlations with disease, opening the door to discovery of clinical covariates and, ultimately, precision medicine applications.

We therefore developed a targeted sequencing panel focused on chronic liver disease driver genes, reasoning that variants under positive selection in chronic livers are most likely to reflect disease biology and offer the greatest potential for clinical utility. We collated 17 established chronic liver disease driver genes from previous liver sequencing studies^15,16^ (Brzozowska et al., manuscript in preparation), which we call chronic liver disease-associated somatic mutation (CASM) genes (Methods; **Supplementary Table 4**). To provide additional context for interpreting cell-free DNA signal, we included genes highly expressed in the liver as tissue-of-origin markers^15,48^ (n = 9), as well as control gene sets from non-chronic liver disease contexts: clonal hematopoiesis drivers^6,49^ (n = 4), hepatocellular carcinoma drivers^50^ (n = 4), inflammatory bowel disease drivers^21–23^ (n = 11), and colorectal cancer drivers^51^ (n = 4) (**Fig. 2A**; **Supplementary Table 4**). The resulting 59-gene panel covered 203 kb with 3× probe tiling and incorporates duplex adapters for molecular error suppression, enabling detection of ultra-low frequency variants at a target raw amplicon coverage of 80,000x (Methods).

**Figure 2:**
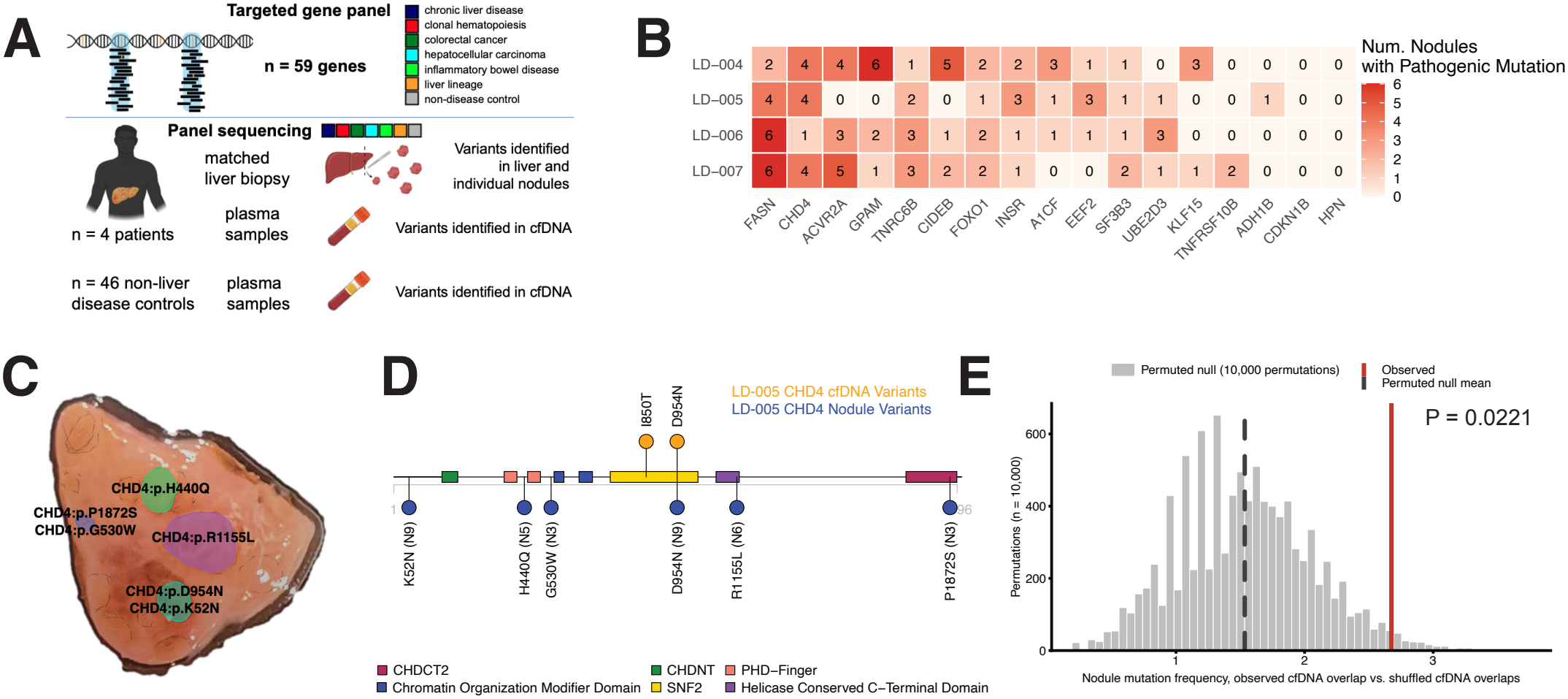
Liver disease-specific panel of CASM genes enables biopsy-naïve non-invasive detection of liver disease-associated somatic variants from distinct liver nodules in cell-free DNA. **A**, Top, targeted panel approach to detect driver gene variants for 59 genes, capturing chronic liver disease-associated somatic mutation (CASM) genes, clonal hematopoiesis driver genes, colorectal cancer driver genes, hepatocellular carcinoma driver genes, inflammatory bowel disease driver genes, genes that are highly expressed in hepatocytes and considered liver marker genes, and non-disease specific control genes. Full information about the panel can be found in Supplementary Table 4. Bottom, schematic of liver disease cohort comprising n = 4 patients with matched liver biopsy, including panel sequencing of 8-10 nodules per patient and cell-free DNA samples, to establish detection of driver variants from livers versus cell-free DNA using the targeted panel. Cell-free DNA from plasma samples from n = 46 non-liver disease controls also underwent targeted panel sequencing. **B**, Heatmap displaying the number of liver nodules carrying an independent pathogenic somatic variant in the indicated CASM genes, from panel sequencing of liver nodule samples for each patient. A variant was counted as independent only if it was found in a single nodule of that patient. Exact variants present in two or more nodules of the same patient were excluded as inherited from a shared ancestor, with the exception of the *FOXO1p*.*S22W* hotspot. **C**, Image of liver segment from patient LD-005 with nodules that were sampled for targeted panel sequencing indicated. Nodules are assigned a number and identified CHD4 variants are annotated in the nodule from which they were identified. **D**, Illustration of the CHD4 protein with indicated functional domains (CHDNT, CHD4 N-terminal domain; PHD-Finger, plant-homeodomain finger; Chromo, chromodomain; SNF2, SNF2-like ATPase motor domain; CHDCT2, CHD C-terminal 2 domain). Somatic CHD4 variants identified from the cell-free DNA are displayed in yellow (top) and variants identified from the liver nodule sequencing are displayed in blue (bottom) for cryptogenic liver cirrhosis patient LD-005. The number assigned to the nodule from which the variant was detected is indicated parenthetically. **E**, Association between cell-free DNA detection of CASM genes and the degree of clonal convergence in matched liver nodule biopsies, assessed across all four liver cirrhosis patients. Each CASM gene in each patient was given a weight equal to the fraction of that patient’s nodules carrying an independently acquired disease-specific mutation in that gene, so a gene mutated in half of a patient’s nodules scores 0.5 and a gene mutated in all of them scores 1. *FOXO1* was restricted to the p.S22W hotspot. The observed score, shown as the red line, is the sum of these weights over the genes detected in that patient’s cell-free DNA, summed across all four patients. Gray bars show the null distribution. In each of 10,000 permutations, every patient was assigned the same number of genes as their cell-free DNA detected, drawn at random without replacement from the full 17-gene CASM panel, and the weights of the drawn genes were summed. The dashed line is the mean of this null. The permutation test breaks only the pairing between genes detected in cell-free DNA and their nodule weights, holding fixed the number of cell-free DNA detections per patient, the 17-gene CASM panel, and the weights themselves. The P value is the fraction of permutations with a score at or above the observed value, calculated as (1 + the number of simulated trials at or above observed) / (1 + 10,000).

To call somatic variants with high specificity in cell-free DNA, we developed a variant calling pipeline integrating duplex denoising based on read-level quality features and an empirical Bayes beta-binomial framework for background error modeling, building on approaches for suppressing stereotypical sequencing errors^38^ and enabling us to call mutations with single-read duplex support (**Supplementary Fig. 5A**, Methods). To assess the specificity of our calling pipeline and panel design for chronic liver disease-relevant somatic evolution, we applied this approach to 36 liver nodules (8 – 10 per patient) and matched plasma cell-free DNA from four cirrhosis patients (LD-004 through LD-007), all with prior whole-genome sequencing (5 of the 41 whole-genome sequenced nodules had insufficient material for panel sequencing) (**Fig. 2A; Supplementary Table 5**). Across 36 nodules, deep duplex sequencing detected large numbers of non-synonymous variants distributed across nearly every gene in the panel, including genes with no established role in MASLD (**Supplementary Fig. 5B**). Many of these likely represent real mutations present at low cellular fractions, reflecting the sensitivity of duplex sequencing; however, their presence does not imply positive selection or contribution to clonal outgrowth. For example, *APOB*, which is not a chronic liver disease driver, harbored the highest mutation burden, and control genes accumulated variant counts comparable to several chronic liver disease driver genes (**Supplementary Fig. 5B**).

We reasoned that pathogenicity scores could distinguish mutations under disease-associated selection from incidental variants accumulated through neutral processes. Analysis of published driver mutation catalogs across chronic liver disease^15^, IBD^22,23^, hepatocellular carcinoma, and colorectal cancer (PCAWG)^50^ confirmed strong enrichment for high pathogenicity scores among established drivers (**Supplementary Fig. 5C**; Methods). Across the four liver samples, *FOXO1, KLF15, GPAM, ACVR2A* and *CIDEB* showed significant enrichment for pathogenic variants (**Supplementary Fig. 5D**). Control genes (IBD, colorectal cancer, and hepatocellular carcinoma genes), despite comparable or greater mutation burden, mostly fell below the significance threshold, with notable exceptions such as *DNMT3A* and *ASXL1*, consistent with clonal hematopoiesis detected in tissue through immune cell infiltration^53,54^. *ARID1A* also had a significant enrichment for pathogenic variants; we did not assign this as a CASM gene because *ARID1A* is also somatically mutated across other tissues and disease contexts^22,23,55^, not exclusively in chronic liver-disease contexts. Liver lineage genes such as *ALB* and *APOB* similarly lacked pathogenic enrichment, consistent with accumulation through a lineage-specific mutational process rather than selection^56^. Aggregating by gene-set category confirmed this pattern: both CASM genes and clonal hematopoiesis genes showed significant pathogenicity enrichment as a group, while the remaining genesets were not significantly enriched for pathogenic variants in the liver (**Supplementary Fig. 5E**).

Convergent somatic evolution, where independent mutations arise in the same driver gene across distinct clones, has been reported in chronic liver disease; previous WGS studies profiling the liver with LCM have revealed chronic livers with up to 14 independent *CIDEB* clones on one segment^15^. Across our four patients with liver cirrhosis and matched plasma cell-free DNA, we identified 18 gene– patient combinations in which three or more independent pathogenic mutations in the same CASM gene arose in different nodules (**Fig. 2B**), with different patients showing selection for different drivers, consistent with distinct metabolic environments^15^. In one striking case, six distinct pathogenic variants in *CHD4* were identified across four nodules in patient LD-005 (**Fig. 2C**); one of these pathogenic variants (p.D954N) was detected with denoised duplex support in plasma cell-free DNA, alongside an additional pathogenic *CHD4* variant not observed in any biopsied nodule (**Fig. 2D**), illustrating how profiling with plasma cell-free DNA can capture somatic clones missed by limited tissue sampling. We also identified a phased *KLF15* variant in patient LD-005, where two pathogenic mutations are contained within a single cell-free DNA fragment^57^, supported by multiple duplex reads in plasma and present in the genomic DNA with duplex support. Notably, cell-free DNA fragments carrying CASM gene variants were shorter than fragments that were wild type at the same loci across all four patients (P = 1.66 x 10^-2^; **Supplementary Fig. 6**), consistent with liver-derived origin. To test whether cell-free DNA detection is biased toward genes under convergent selection, we weighted each CASM gene by the fraction of that patient’s nodules carrying an independently acquired pathogenic mutation, summed these weights over cell-free DNA-detected genes across all four patients, and compared the observed statistic to a permutation null (10,000 iterations; **Methods**). Cell-free DNA detections were significantly enriched for convergently mutated genes (P = 0.0221; **Fig. 2E**), demonstrating that our panel preferentially captures signal from CASM genes under active selection (i.e., showing evolutionary convergence) within individual livers, compared to genes that do not show convergent evolution.

Collectively, these results establish that our targeted panel non-invasively detects cell-free DNA derived from chronic liver disease clones, with specificity enhanced by pathogenicity filtering and detection biased toward CASM genes under convergent selection, paving the way for application to larger cohorts without matched liver biopsies.

### Non-invasive monitoring of chronic liver disease driver variants using deep, targeted sequencing in large cohorts

We next expanded our analysis to a larger cohort of chronic liver disease patients with plasma cell-free DNA but without matched liver biopsies. In addition to the four aforementioned patients with matched explants, this cohort comprised 147 chronic liver disease patients sequenced across two independent cohorts: 72 patients with MASLD and alcohol-related liver disease from the DeLIVER consortium (sequenced at NYGC), and 75 patients with MASLD, alcohol-related, and other cirrhosis etiologies from UT Southwestern (sequenced at UTSW), analyzed together with the 46 non-liver-disease control plasma samples (**Fig. 3A**; **Supplementary Table 1**). All plasma samples were sequenced with our targeted panel and duplex DNA error correction on the Illumina platform (Methods; **Supplementary Table 6**).

**Figure 3:**
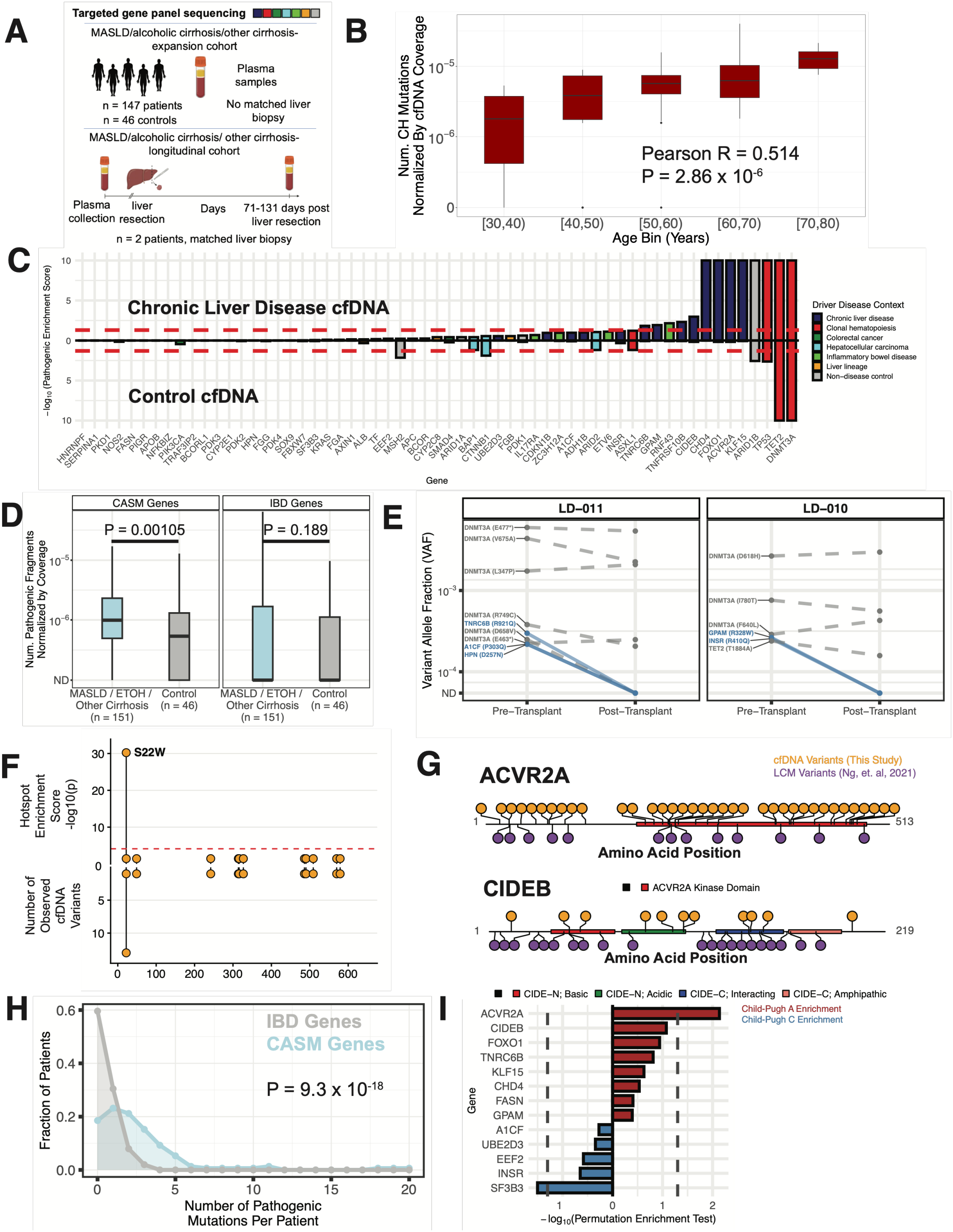
Non-invasive monitoring of CASM gene variants from targeted panel sequencing allows for patient comparisons and identification of clinical covariates of cell-free DNA from large cohorts. **A**, Schematic of expansion cohort for biopsy-naïve detection of CASM gene variants from plasma only (top). The targeted panel was also applied to a longitudinal cohort comprising two patients with plasma sampling before and after liver resection surgery (bottom). Targeted sequencing with duplex error correction was performed with Illumina. **B**, Number of pathogenic variants in clonal hematopoiesis (CH) panel driver genes normalized by library size from the cell-free DNA across patient age. The bottom and top ends of the boxes represent the 25th and 75th percentiles of the data, respectively, and the horizontal lines represent the median. The whiskers represent at most 1.5 times the interquartile range. Two-sided t-test. N = 74 patients with available age information were included in this analysis. **C**, Pathogenicity enrichment scores for driver genes associated with specific disease contexts calculated from somatic variants detected in cell-free DNA for the expanded cohort (n = 151 chronic liver disease patients; top) or non-liver disease controls (n = 46; bottom). For each gene, the total number of somatic variants observed are tabulated and each variant is assigned an AlphaMissense score^67^ and a CADD Raw score^68^. The null set for each gene is every panel-interrogated site in that gene, with each site weighted by the background error rate at that site, so that positions more likely to generate a call are drawn more often. The same number of mutations as observed in that gene is drawn from this null with replacement, and the mean AlphaMissense and mean CADD Raw scores of the drawn set are calculated. The sampling is repeated 10,000 times per gene. The *P* value is the fraction of draws in which the null mean exceeds the observed mean. AlphaMissense and CADD *P* values are calculated separately and then combined with Fisher’s method for combining *P* values. *P* values were corrected across genes with the Benjamini-Hochberg method. The red dashed line represents the significance threshold of 0.05. **D**, Detection of SNVs in CASM genes (left) or IBD driver genes (right) in cell-free DNA using the targeted sequencing panel for non-liver disease controls (n = 46 plasma samples) versus chronic liver disease patients (n = 151 plasma samples) without matched liver biopsy. Genes were assigned to one specific group (CASM or IBD). For each group, the total number of fragments with base quality >30 supporting a denoised variant is divided by the total number of duplex fragments sequenced across genes belonging to the respective group. The bottom and top ends of the boxes represent the 25th and 75th percentiles of the data, respectively, and the horizontal lines represent the median. The whiskers represent at most 1.5 times the interquartile range. Two-sided Wilcoxon test. ns, not significant. **E**, Longitudinal analysis of chronic liver disease-associated variants detected in the cell-free DNA from two patients with samples before and after liver transplant. The number of pathogenic variants normalized by coverage is plotted at both time points. Variants in CASM genes (blue) that were detected at the pre-transplant timepoint are undetectable after transplant. Variants in clonal hematopoiesis-associated driver genes *DNMT3A* and *TET2* (grey) that were detected at the pre-transplant timepoint maintain comparable levels after transplant. **F**, Hotspot enrichment score for variants detected in *FOXO1* showing enrichment at the p.S22W hotspot (upper plot). To obtain the hotspot enrichment score, we first counted the number of unique observations per amino acid change in FOXO1. We then calculated the total number of observed mutations. The null probability of observing a mutation at any single site was defined as the inverse of the number of FOXO1 codons covered by the sequencing panel (n = 656), so that the expected distribution is determined by the interrogated region rather than by the observed mutations themselves. We then used a one-sided binomial test to test whether each observed count of a specific amino acid change exceeded the expected count under this null probability, identifying potential hotspots with significantly higher than expected recurrence. Because every covered codon is implicitly tested with deep targeted duplex sequencing, and not only those at which a mutation was observed, the multiple testing family was defined as all 656 covered codons rather than the smaller number of codons carrying a detected variant. The red dashed line indicates the Bonferroni-corrected significance threshold for this family (α = 0.05/656 = 7.6 × 10^−5^, plotted at −log_10_P = 4.12); only p.S22W exceeds this threshold. Bottom plot displays the number of *FOXO1* variant detections in cell-free DNA from 151 chronic liver disease patients. **G**, Annotated ACVR2A (top) and CIDEB (bottom) protein plots displaying cell-free DNA variants detected in the current study (yellow, top) and liver variants identified from whole-genome sequencing of laser capture microdissections from Ng et al.^15^ (purple, bottom). ACVR2A and CIDEB domains are indicated. **H**, Number of pathogenic mutations per patient in chronic liver disease CASM genes (light blue, 17 genes) and IBD genes (grey, 11 genes) for the same n = 151 chronic liver disease patients identified from panel sequencing of the cell-free DNA. *FOXO1* mutations were restricted to S22W. Points show the fraction of patients carrying the indicated number of mutations; patients with 0 pathogenic mutations are the leftmost point of each curve. The x axis is truncated at 20. Counts are paired within patient. Patients have a median 2 more pathogenic mutations in CASM genes versus IBD driver genes, P = 9.3 x 10^-18^, one-sided Wilcoxon signed-rank test. **I**, Permutation enrichment test of liver disease severity for samples carrying pathogenic variants in the indicated CASM genes. For each gene, the observed statistic was the mean expected Child-Pugh score, calculated by assigning the midpoint of each Child-Pugh class (A = 5.5, B = 8, C = 12.5) to each mutation-positive sample. This value was compared with a null distribution generated from 10,000 permutations by randomly sampling the same number of patients from the overall cohort-wide Child-Pugh class distribution (71 Child-Pugh A, 43 Child-Pugh B, and 31 Child-Pugh C, 6 samples were excluded due to missing Child-Pugh status). Bars represent the signed permutation enrichment statistic. The dotted grey lines denote the nominal significance threshold (two-sided permutation P = 0.05).

As a positive control, we first analyzed cell-free DNA variants detected in clonal hematopoiesis genes across patient age^6,7,58,59^. As expected, we identified increasing variant fractions with age (**Fig. 3B**). We then applied the pathogenicity enrichment across all 151 chronic liver disease and 46 control plasma samples (**Fig. 3C**). Clonal hematopoiesis genes showed enriched pathogenicity in both liver patients and controls, consistent with the near-ubiquitous presence of clonal hematopoiesis variants in adult blood^6,7,58^. In contrast, CASM genes showed significant pathogenicity enrichment only in the liver disease cohort, where 8 CASM genes reached significance.

Plasma samples from the chronic liver disease cohort had significantly higher numbers of pathogenic mutations in CASM genes compared to non-liver disease controls (P = 1.05 x 10^-3^; **Fig. 3D**). In contrast, there was no significant difference in the number of detected pathogenic variants in IBD-associated genes between the liver-disease vs the control cohorts (P = 0.189; **Fig. 3D**, see **Supplementary Fig. 7A** for similar comparison without filtering for pathogenic variants). Stratifying by sequencing center, the chronic liver disease driver gene signal replicated independently in both the DeLIVER cohort (sequenced at NYGC) and the UTSW cohort, with no significant difference in IBD-associated genes **(Supplementary Fig. 8)**, confirming that findings are robust across sites. Unlike clonal hematopoiesis, CASM gene detection did not correlate with age (**Supplementary Fig. 9**), providing further evidence that the chronic liver disease cell-free DNA signal is distinct from clonal hematopoiesis and reflects liver-derived biology. We additionally note that 9 of our 17 CASM genes were observed in previous LCM sequence files from MASLD and alcohol-related liver disease etiologies^15,60^: *FOXO1p*.*S22W, CIDEB, GPAM, ACVR2A, TNRC6B, CHD4, INSR, FASN*, and *A1CF*.

Restricting to this set, we again observed statistically significant detection of pathogenic variants in chronic liver disease plasma (**Supplementary Fig. 7B**). Detection was significantly higher in MASLD/alcohol-related liver disease than in other cirrhosis etiologies (primary biliary cholangitis, hepatitis B, hepatitis C; **Supplementary Fig. 7C**), consistent with selection of these drivers by the metabolic environment. This argues the cell-free DNA signal reflects metabolic-specific selective pressure rather than generic hepatocyte turnover from liver injury.

To explore the ability to serially monitor pathogenic CASM gene variants, we profiled pre- and post-transplant plasma samples from two patients who received liver transplantation (**Fig. 3A**, bottom). In the pre-transplant plasma, we detected fragments with pathogenic variants in at least one CASM gene in both patients, including three genes in LD-011 (**Fig. 3E, Supplementary Fig. 10**), while fragments with pathogenic variants were undetected at the post-transplant time point. In contrast, the fragments with pathogenic variants in the clonal hematopoiesis genes *DNMT3A* or *TET2* remained at similar levels at the two time points across patients, consistent with a non-liver origin.

To assess whether cell-free DNA variants reflected known patterns of functional selection in CASM genes, we mapped the detected variants in our cohort of 151 chronic liver disease patients to protein domain structures and compared their localization to previous liver biopsy sequencing data^15^ (**Fig. 3F**,**G, Supplementary Fig. 11**). Specifically, there were eight genes included in our panel that had variants observed in the original study profiling chronic liver disease genomes through laser capture microdissection (*FOXO1, ACVR2A, CIDEB, INSR, GPAM, CHD4, FASN*, and *TNRC6B*)^15^. For *FOXO1*, we detected the recurrent gain-of-function hotspot p.S22W, which disrupts AMPK-mediated phosphorylation and promotes nuclear retention of FOXO1, conferring a selective advantage in the steatotic liver environment^15^. Indeed, only the p.S22W hotspot exhibited a recurrent signal, observed across multiple cell-free DNA samples in our cohort (**Fig. 3F**). Similarly, the majority of our detected *ACVR2A* variants fell within the kinase domain (**Fig. 3G**, top, **Supplementary Fig. 12A**), consistent with previous reports^15^. *CIDEB* cell-free DNA variants impacted domains essential for lipid droplet fusion^61^ (**Fig. 3G**, bottom). *INSR* variants were enriched in the tyrosine kinase domain and exhibited the same 3’ clustering reported in laser capture microdissection data (**Supplementary Fig. 11A**). *GPAM* variants affected the acyltransferase domain (**Supplementary Fig. 11B, Supplementary Fig. 12B**), *CHD4* variants again targeted the SNF2 motor domain (**Supplementary Fig. 11C**), *FASN* variants fell within the beta-ketoacyl synthase, acyltransferase, and ketoacyl reductase domains (**Supplementary Fig. 11D**)^62–64^, and *TNRC6B* variants fell within the C-terminal domain responsible for mRNA silencing^65^ (**Supplementary Fig. 11E**). Overall, we detected pathogenic variants in CASM genes in the cell-free DNA in 81.5% (123/151) of patients, with a median of two CASM gene variants per patient (range 1-19) (**Fig. 3H**). Patients had a median 2 more CASM gene variants compared to IBD gene variants detected in cell-free DNA (P = 9.3 x 10^-18^; **Fig. 3H**), with a median of 6 reads per CASM geneset per sample and median of 0 reads per IBD geneset per sample (Wilcoxon signed-rank test, one-sided, paired P = 1.44 x 10^-15^). Thus, using our targeted panel, we are able to recover CASM gene variants across the majority of our cohort, and these mutations are found at the expected hotspot location or within functional domains, again providing support that the cell-free DNA fragments bearing these variants are derived from diseased-liver clones.

Taking advantage of our cohort size and available clinical metadata (**Supplementary Table 6**), we next asked whether cell-free DNA CASM gene variants are associated with liver disease severity. We stratified patients by Child-Pugh class [A (mild), B (moderate), and C (severe hepatic impairment)^66^] and computed enrichment scores for chronic liver disease driver genes with at least 5 variants across patients. We identified differential enrichment for CASM genes, with *ACVR2A* pathogenic variants being significantly enriched in Child-Pugh A patients (**Fig. 3I**). This underscores that different mutant clones may have distinct effects depending on disease stage, potentially pointing to changing selective pressures that shape somatic evolution in diseased livers.

### Non-invasive detection of IBD driver mutations in cell-free DNA

Having established that our targeted duplex sequencing panel combined with pathogenicity-based enrichment can detect disease-associated somatic signal in the plasma of patients with chronic liver disease, we next asked whether the same framework generalizes to a second, mechanistically distinct disease of the gastrointestinal tract: IBD. Somatic mutations in a recurrent set of driver genes, including *NFKBIZ, PIGR, TRAF3IP2*, and *ZC3H12A*, have been previously identified in the inflamed colonic epithelium of ulcerative colitis (UC) patients^22,23^, and our custom panel was designed to include these loci alongside the CASM set. We applied our identical read-level denoising, empirical Bayes background error modeling, and pathogenicity-based enrichment pipeline to cell-free DNA from 10 patients with biopsy-confirmed active UC inflammation and 14 patients with biopsy-confirmed prior inflammation, and benchmarked detection against our non-liver disease, non-IBD control cohort (n = 46; **Fig. 4A; Supplementary Table 7**).

**Figure 4:**
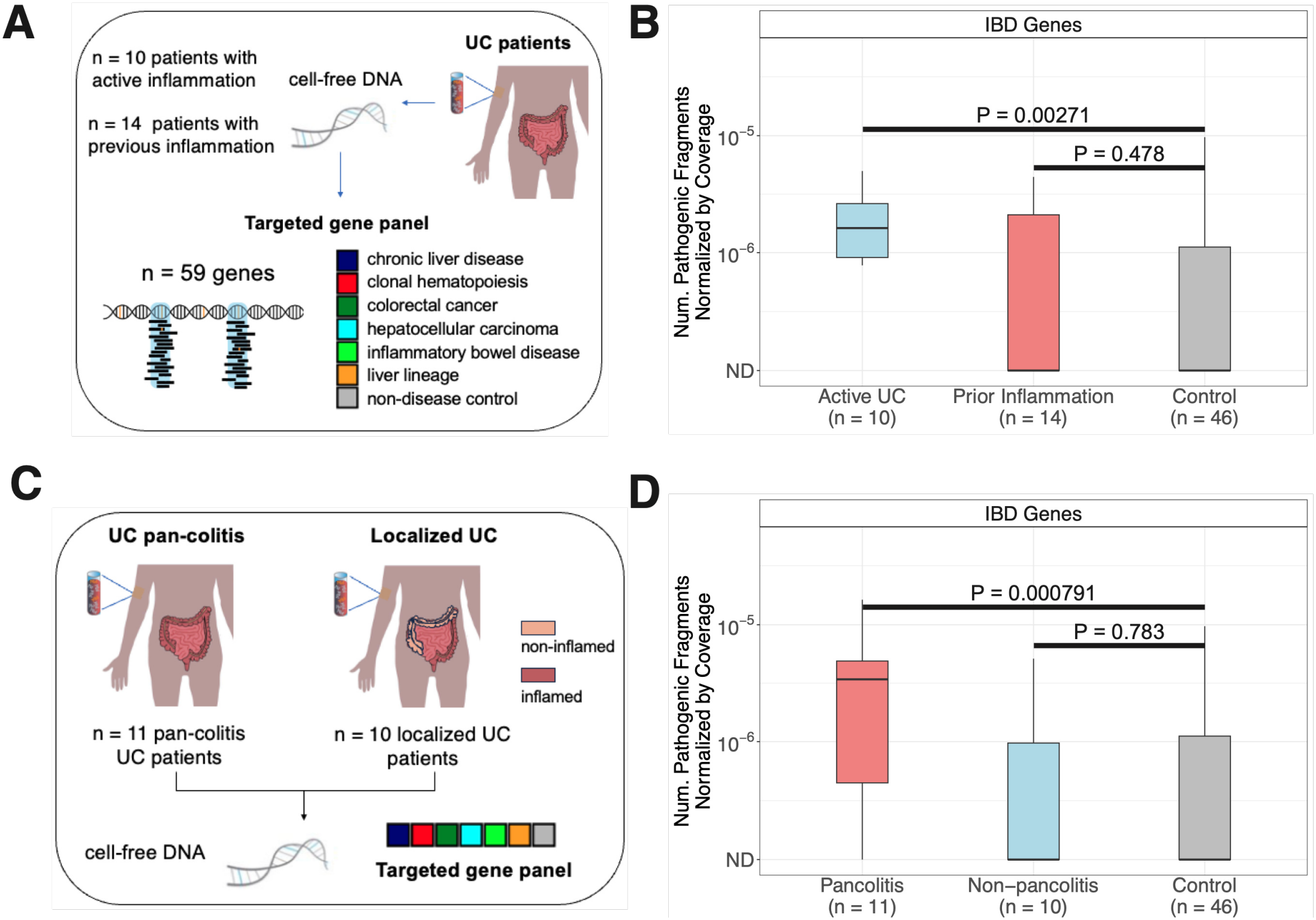
Non-invasive monitoring of IBD SNVs and indels using targeted panel sequencing. **A**, Study schematic. Plasma cell-free DNA was collected from patients with inflammatory bowel disease (IBD), specifically ulcerative colitis (UC), including individuals with active inflammation (n = 10) and those with a previous history of inflammation (n = 14). Plasma samples were subjected to targeted sequencing using the identical custom driver gene panel employed across the study to identify mutations in known IBD driver genes. Pathogenic enrichment analysis was applied to detect recurrent hotspots and functionally impactful driver mutations from plasma-derived variants. **B**, IBD driver mutation detection rates. Boxplots depict the number of pathogenic fragments, classified using AlphaMissense and CADD scores, normalized by cell-free DNA coverage in IBD driver genes for UC patients with active inflammation, prior inflammation and non-liver disease controls. ND indicates not detected. Boxes represent the interquartile range (25th–75th percentiles), center lines indicate the median, and whiskers extend to 1.5× the interquartile range. *P* values were calculated using two-sided Wilcoxon rank-sum tests comparing active UC IBD group to controls. **C**, Miami Plasma cell-free DNA IBD Study schematic. Plasma cell-free DNA was collected from patients with inflammatory bowel disease (IBD; specifically ulcerative colitis), including individuals with pancolitis (n = 11) and those with localized inflammation (n = 10; inflammation restricted to left-side, ascending, rectum, recto-sigmoid or sigmoid-cecum). Plasma samples were subjected to targeted sequencing using the identical custom driver gene panel employed across the study to identify mutations in known IBD driver genes. Low depth samples (less than 200x simplex coverage) were removed from this cohort and corresponding controls for this analysis. **D**, IBD driver mutation detection rates. Boxplots depict the number of pathogenic fragments, classified using AlphaMissense and CADD scores, normalized by cell-free DNA coverage in IBD driver genes for UC patients with pancolitis, non-pancolitis (localized inflammation; Supplementary Table 7) and non-liver disease controls. ND indicates not detected. Boxes represent the interquartile range (25th–75th percentiles), center lines indicate the median, and whiskers extend to 1.5× the interquartile range. *P* values were calculated using two-sided Wilcoxon rank-sum tests comparing pancolitis UC IBD group to controls.

Patients with active UC showed elevated detection of pathogenic variants in IBD driver genes compared to controls (P = 2.71 x 10^-3^, **Fig. 4B**), consistent with the hypothesis that active inflammatory flares shed clonally expanded intestinal epithelial DNA into circulation at levels sufficient for detection with our panel. In contrast, UC patients without actively inflamed biopsies (n = 14) did not show significant elevation in IBD driver gene detection compared to controls (P = 0.478, **Fig. 4B**), suggesting that active inflammation is required to generate sufficient epithelial turnover for cell-free DNA signal.

For further validation of this tool, we assessed a second UC cohort comprising individuals with pancolitis (n = 11) and those without pan-colitis (n = 10; inflammation restricted to left-side, ascending, rectum, recto-sigmoid or sigmoid-cecum, which we call localized inflammation; **Fig. 4C, Supplementary Table 7**). Here, pancolitis impacts the entire colorectal tissue, while localized inflammation is spatially restricted to smaller regions. We applied the targeted panel to plasma cell-free DNA from this cohort, observing that patients with pancolitis had significantly higher detection levels of pathogenic variants in IBD driver genes compared to controls (P = 7.91 x 10^-4^), while patients with localized inflammation did not show significant elevation in IBD driver gene detection compared to controls (P = 0.783, **Fig. 4D**). These results suggest that the disease signal in the cell-free DNA tracks with the tissue area that is affected by inflammation, with more extensive pancolitis leading to more shedding. Together, these results extend our framework beyond chronic liver disease and argue that non-invasive detection of disease-associated somatic variants in cell-free DNA is generalizable across chronic inflammatory diseases with defined driver mutation repertoires.

## Discussion

Somatic mosaicism in non-malignant tissues is rapidly emerging as a discovery engine for human disease biology and a complementary axis to germline genetics for therapeutic target identification^69^. Where germline variation identifies genes and pathways causally implicated in disease across populations, revealing what is tolerable across development, somatic variation reveals what is selectively advantageous within a specific diseased tissue environment — a natural, in-human genetic screen running across decades of pathology, in millions of cells, under the precise selective pressures we hope to therapeutically modify. Recent work has shown that this somatic substrate is unusually rich in actionable biology: loss-of-function mutations in *CIDEB* and *GPAM* in MASLD livers nominate lipid-metabolism vulnerabilities, now in active clinical development (CIDEB-siRNA, NCT06836609). IL-17 pathway mutations in inflamed IBD epithelium predicted, correctly, that IL-17 inhibition would worsen rather than ameliorate colitis^21–23^. *SERPINA1* truncation variants in α1-antitrypsin deficiency reveal a novel therapeutic principle^70^. Beyond target discovery, the patient-specific clonal landscape itself carries information: MASLD patients with *FOXO1*-dominant, *GPAM*-dominant, or *CIDEB*-dominant clonal profiles may be responding to genuinely distinct upstream pressures (e.g., excess carbohydrate vs. excess dietary fat), suggesting that the somatic mutational composition of a diseased tissue could function as a companion diagnostic, stratifying patients within a clinical category into mechanistically distinct subgroups and pointing toward distinct therapies.

Realizing this potential, however, has been blocked by a major barrier: tissue access. Tissue biopsy is rarely clinically indicated in chronic disease or carries morbidity such as annual colonoscopies in long-standing UC. While surgical explants and rapid autopsies have produced the field’s foundational datasets, these cannot scale. Moreover, the spatial organization of solid tissues means that even when tissue is available, a single biopsy captures only a small patch of a much larger mosaic. This sampling bottleneck has restricted the field to small cross-sectional studies, foreclosed longitudinal analyses, and prevented the kind of large-cohort epidemiology that turned clonal hematopoiesis from an observation into a growing field of translational science.

In this work, we show that this barrier can be lowered. Applying ultra-sensitive cell-free DNA sequencing to patients with chronic liver and bowel disease, we detect clone-resolved somatic signal at fractions comparable to cancers, track it across the spatial and phylogenetic structure of matched tissue, scale it to a 151-patient cohort (147 biopsy-naive, 4 with matched explant), observe its disappearance after curative resection, and connect it to disease stage. The clonal composition of chronically diseased solid tissues can thus be read from the blood. The trajectory this enables is prefigured by clonal hematopoiesis: when CHIP became detectable in peripheral blood, it became, within a decade, a major axis of aging, cardiovascular, metabolic, and cancer research^6,24,25,71,72^. Our results suggest that liquid biopsy may now provide a similar path of growth to the field of somatic genetics in solid tissue chronic disease.

Beyond the methodological advance, our data already begin to surface the biology of solid-tissue CM in chronic disease. The shedding rates we observe in MASLD (10^−4^–10^−5^) are comparable to stage II-III colorectal cancer and above non-invasive precancerous lesions^29^, plausibly reflecting ongoing hepatocyte turnover, disrupted tissue architecture, or inflammatory cell death. Cell-free DNA fragments carrying liver-derived variants are systematically shorter than wild-type fragments, paralleling the fragmentation signature of circulating tumor DNA and suggesting that the chromatin and nucleosomal state of chronically diseased cells may be altered in ways legible from plasma^42^. We further observed marked spatial and phylogenetic heterogeneity in shedding across neighboring liver nodules within a single patient, indicating that clonal shedding is not a uniform function of clonal burden but a clone-intrinsic property shaped by local tissue environment, an observation that motivates future dissection of which features of the diseased microenvironment promote cell-free DNA release. The stage-specific enrichment of *ACVR2A* variants in earlier-stage Child-Pugh A disease hints that the driver gene composition of the diseased liver shifts across disease stages, consistent with evolving selective pressures and raising the possibility that different clones are adaptive at different points in the disease trajectory. Extending the same framework to inflammatory bowel disease, we detect somatic variants in IL-17 pathway genes, the same loci whose loss-of-function in colonic epithelium predicted, in advance of clinical confirmation, that IL-17 inhibition would worsen colitis^22,23^. That this pathway is detectable from cell-free DNA in active UC patients and those with pancolitis suggests that the somatic clonal record of chronic inflammation accumulates in ways accessible to non-invasive surveillance.

Several limitations frame the interpretation of this work. Tissue-of-origin assignment is not definitive: some panel targets are expressed beyond the liver, and somatic driver variants are known to occur in normal tissues outside the organ in which they have been most studied (e.g., *CTNNB1* in normal stomach^55^). Our strongest evidence for liver origin comes from post-transplant signal clearance and spatial concordance with matched biopsies, but absolute tissue-of-origin confirmation in the broader panel cohort remains indirect. Methylation-based deconvolution^27,28^ may address this in future iterations. Our pathogenicity-based enrichment is biased toward loss-of-function variants and may miss gain-of-function or regulatory mutations that contribute to clonal fitness. Importantly, we still detect chronic liver disease-specific signal when restricting to nonsynonymous variants without pathogenicity filtering. Sample-level effect sizes in the expansion cohort, while statistically robust, overlap with controls at the individual-sample level. Refined panels focused on hotspot regions, integration with orthogonal cell-free DNA modalities (fragmentomics, methylation), and larger validation cohorts will be required to bridge from population-scale signal to single-patient clinical assay.

Looking forward, the framework established here defines a future research program along two axes. First, as a tool for discovery science. The somatic mosaicism of diseased tissues is, in effect, a natural in-human genetic screen, where decades of disease-specific selective pressure point toward the genes and pathways that confer fitness in pathological environments^69^. Reading this screen from plasma rather than tissue dramatically expands the scale and resolution at which it can be interrogated. Longitudinal sampling, until now infeasible, can begin to map how the clonal landscape evolves across stages of chronic disease, which clones emerge early, which persist, and which are replaced. Sufficiently powered cohorts can connect specific driver clones to the clinical, environmental, and germline contexts in which they expand, building a more granular picture of how disease shapes somatic evolution and how somatic evolution, in turn, shapes disease. The framework also opens the discovery of new drivers in diseases where access to tissue has been the central limitation. By applying ultra-deep duplex whole-exome sequencing to cell-free DNA, somatic drivers of chronic diseases that are rarely biopsied, including chronic kidney disease, endometriosis, chronic lung disease, and many others, may be identified directly from plasma. Our MASLD shedding rates, comparable to those of cancers, suggest that at least for some chronically diseased solid tissues, cell-free DNA shedding will not be the limiting factor it might have been assumed to be.

Second, as a foundation for precision medicine. Patient-level driver gene composition, accessible from a blood draw, may resolve mechanistically distinct subgroups within clinical categories that are currently treated as homogeneous. Chronic liver disease patients whose plasma reveals predominantly *FOXO1, GPAM*, or *CIDEB*-driven clonal landscapes may reflect genuinely different upstream etiologies and, plausibly, different optimal therapies among the mechanism-specific agents now entering clinical development^69^. Longitudinal cell-free DNA monitoring may then track whether the clonal landscape shifts in response to a chosen therapy, providing a non-invasive readout of treatment response at the level of the cells under selection. The same logic of stratify, treat, and monitor has transformed oncology over the past two decades through the integration of tumor sequencing into routine care. Solid-tissue clonal mosaicism, now accessible from plasma, opens a path to extending that paradigm into chronic non-malignant disease, the somatic genetic frontier that has, until now, been the exclusive province of cancer.

## Acknowledgments

We thank the patients and their families for contributing plasma and tissue for this study. We also thank members of the Landau laboratory for input, as well as Jane Park for outstanding technical support and Henry Rui for operational support. This work was supported by the Mark Foundation Aspire Award, Vallee Scholar Award, Burroughs Wellcome Fund Career Award for Medical Scientists, a National Cancer Institute R01 grant (R01-CA266619-01), the Melanoma Research Alliance Established Investigator Award (D.A.L.). D.A.L. is a Blood Cancer United Scholar. H.Z. is supported by the Simmons Comprehensive Cancer Center (NCI Cancer Center Support Grant P30CA142543 and SCCC UTSW translational pilot grant), Cancer Prevention and Research Institute of Texas (RP220614) and the National Institutes of Health (DK139976). T.R. is supported by the NIHR Cambridge Biomedical Research Centre (grant number BRC-1215-20014) (T.R.) and the Wellcome Trust (grant numbers 206194, 220540/Z/20/A and 108413/A/15/D). E.B. is supported by the Oxford NIHR Biomedical Research Centre and is an NIHR Senior Investigator. The views expressed in this article are those of the author and not necessarily those of the NHS, the NIHR, or the Department of Health. We would like to acknowledge the DeLIVER consortium who supported this study. This work was supported in part by funding from Montage Bio, Inc., under an industry-sponsored research agreement with Cornell University. This work was made possible by the MacMillan Family Foundation and the MacMillan Center for the Study of the Non-Coding Cancer Genome at the New York Genome Center. The opinions, results, and conclusions reported in this paper are those of the authors and are independent from these funding sources.

## Author Contributions

S.R., A.P.C. and D.A.L. conceived and designed the project.

A.Ru., E.W., R.F., I.R. and A.J. performed library preparation and sequencing.

S.R., B.L., A.P.C., J.Z., A.R.D., T.B., W.D., A.Ra., W.S.L., J.P. and A.J.W. performed data analysis.

N.B., F.J.R., M.H. and P.J.C. provided liver biopsy sequencing data and plasma samples from chronic liver disease patients and clinical/biological expertise informing the chronic liver disease results.

M.Z., A.A., A.Sh., M.W., D.H., A.Si. and M.P. provided liver biopsy and plasma samples from chronic liver disease patients.

H.Z. provided liver biopsy and plasma samples from chronic liver disease patients and clinical/biological expertise informing the chronic liver disease results.

E.A., J.R. and E.B. provided plasma samples from chronic liver disease patients and clinical/biological expertise informing the chronic liver disease results.

M.T.I.B., C.C.M., M.P., W.G., T.R., M.T.A. and C.A.A. provided plasma samples from IBD patients and clinical/ biological expertise informing the IBD results.

S.R., A.P.C., C.P. and D.A.L. wrote the manuscript with comments and contributions from all authors.

## Code and Data Availability

Code and custom scripts will be made available as a GitHub repository.

The raw genomic sequencing data will be deposited to the European Genome-Phenome Archive.

## Methods

### Human samples

All human samples included in this study were collected with informed patient consent, in accordance with ethical guidelines and institutional regulations.

For Wellcome Sanger Institute liver and serum samples, samples from n = 3 MASLD patients were collected with informed consent from Addenbrooke’s Hospital, Cambridge, UK, according to procedures approved by the East of England Local Research Ethics Committee (16/NI/ 0196 and 15/EE/0351). Liver sequencing was performed in^15^ and VCF files from liver biopsies were used for analysis in this current study. Serum samples were collected from the same patients and WGS was performed at NYGC.

For UTSW samples from chronic liver disease patients, the UT-Southwestern Institutional Review Board (IRB) approval for the collection and analysis of tissue samples was granted under protocol #STU 062013-063, while blood sample collection was approved under protocol #STU 092013-010. Clinical plasma samples were obtained from 79 chronic liver disease patients (4 with matched explant nodules and 75 without matched liver biopsies) and 41 non-chronic liver disease controls. The samples were collected and processed uniformly as described below, minimizing pre-analytical variability. For UTSW liver biopsy samples (n = 4), liver tissue samples were collected from patients undergoing liver transplantation. Patient information can be found in **Supplementary Table 1**. WGS (for n = 4 patients) and targeted panel sequencing were performed at NYGC. WGS control samples are from^29^. Five additional control samples from the NYGC WGS control cohort with remaining cell-free DNA material were also sequenced on the targeted panel, giving 46 non-liver disease controls in total (**Supplementary Table 6**).

Approval for the DeLIVER study (PEARL - Prospective Cohort for Early Detection of Liver Cancer^73^) has been obtained from NHS/HRA Research Ethics Committee: East Midlands - Leicester South Research Ethics Committee (REC reference: 21/EM/0151, IRAS 285362), and this study has been registered on ClinicalTrials.gov as NCT05541601.

For plasma samples from IBD patients, patients with UC undergoing routine surveillance colonoscopy were enrolled at Addenbrooke’s Hospital, Cambridge, UK, under NHS Health Research Authority ethics approval (East of England – Cambridge South Research Ethics Committee, REC 17/ EE/0338), with written informed consent obtained from all participants. Inflammation status was assigned by the endoscopist at the time of biopsy, verified against prior colonoscopy and pathology reports in National Health Service (NHS) records, and histologically confirmed by review of H&E-stained sections of the biopsied tissue. Patient-level clinical data were obtained from NHS electronic health records. All patient information can be found in **Supplementary Table 7**. 10 ml peripheral blood was collected in EDTA tubes concurrently for plasma isolation. For plasma samples from pancolitis and non-pancolitis samples, UC patients were enrolled as part of ongoing biospecimen collection at the University of Miami’s Crohn’s and Colitis Center; all subjects provided informed consent (Institutional Review Board protocol 20081100). Intestinal inflammation was assessed by endoscopy and patients were classified as having pancolitis or non-pancolitis. Patient information can be found in **Supplementary Table 7**. Blood samples were collected at the time of colonoscopy. For all plasma samples from IBD patients, targeted panel sequencing was performed at NYGC.

### Sample processing

UTSW: Blood samples were drawn into Streck tubes, which are specifically designed to stabilize cell-free DNA and prevent genomic DNA release. The samples were gently inverted several times to ensure proper mixing with the preservative agents. To isolate plasma, blood tubes were centrifuged at 3000 rpm for 10 minutes at 4°C. The supernatant plasma was carefully transferred to clean RNase/DNase-free tubes, taking care to avoid disturbing the buffy coat components. Plasma aliquots were immediately stored at -80°C until further processing. For the liver biopsy samples from n = 4 patients, fresh tissue was placed into 50 mL conical tubes containing 20 mL of DMEM with 10% DMSO and kept on ice during transport to the laboratory. Upon arrival, the culture medium was discarded, and the tissue was immediately stored at -80°C until further processing. For DNA extraction, tissue samples were thawed on ice, and approximately 30 mg of tissue was sliced into 1–3 mm thick sections using a clean scalpel blade. The tissue slices were transferred to a 1.5 mL microcentrifuge tube and kept on dry ice. Genomic DNA was extracted from liver tissues according to the protocol in^74^. To ensure high DNA integrity, tissues were manually homogenized with a pestle, followed by DNA extraction using the QIAshredder and QIAGEN AllPrep DNA/RNA Mini Kit according to the manufacturer’s protocol.

Extracted genomic DNA was stored at -80°C until further analysis.

DeLiver samples were processed as in^73^.

Wellcome Sanger Institute (IBD Samples): Plasma was isolated from whole blood (∼10 mL) by two sequential centrifugations at 1600 × g (4 °C, brake off; 15 min then 10 min), with the plasma transferred to a fresh tube between spins to deplete residual cells and platelets. Aliquots were stored in 2 mL low-binding tubes at –80 °C prior to downstream processing.

### Illumina WGS cell-free DNA Preprocessing

Samples undergoing WGS were sequenced on an Illumina NovaseqX using 2×150bp read lengths. Samples were demultiplexed using Illumina’s bcl2fastq (v.2.17.1.14) to generate FASTQ files. For cell-free DNA, we used a modified alignment pipeline to accommodate adaptor trimming after observing increased adaptor contaminated reads in cell-free DNA samples compared to tumor samples, given that cell-free DNA molecules can be shorter than the read length (here, 150bp). We therefore used Skewer for adaptor trimming (default settings) and subsequently aligned samples using BWA-MEM (default settings) to the GRCh38 reference (1000 Genomes version). For all cell-free DNA samples, duplicate marking and sorting was conducted using NovoSort MarkDuplicates (v.3.08.02), a multi-threaded bam sort/merge tool by Novocraft Technologies (http://www.novocraft.com), followed by base quality score recalibration using GATK (v.4.1.8; https://software.broadinstitute.org/gatk), resulting in a final coordinate sorted bam file per sample. Alignment quality metrics were computed using Picard (v.2.23.6; QualityScoreDistribution, MeanQualityByCycle, C o l l e c t B a s e D i s t r i b u t i o n B y C y c l e, C o l l e c t A l i g n m e n t S u m m a r y M e t r i c s, CollectInsertSizeMetrics and CollectGcBiasMetrics) and GATK (v.4.1.8; average coverage, percentage of mapped and duplicate reads).

### Biopsy-Informed cell-free DNA SNV Detection in LCM-Profiled Livers (Illumina)

Cell-free DNA reads at patient-specific somatic variant sites were denoised against a biopsy-informed variant set. Per-nodule WGS variant calls were unioned across nodules to give the patient-wide somatic VCF.

Variant sites were first filtered at the locus level. Sites falling in centromeres, simple repeats, or the ENCODE blacklist regions were removed^29^. Variants recurring across our plasma WGS database^29^ above threshold of two appearances were removed, as were variants recurring above threshold of 4 across our MASLD liver cell-free DNA cohort.

The patient-specific set was restricted to variants observed in exactly one biopsy, which removes suspected clonal hematopoiesis and germline variants shared across biopsies. Sites with a Umap k100 mappability score below 1 were removed from both the called set and the coverage denominator. The Umap and Bismap tracks score each position by the uniqueness of the overlapping k-mers, so a k100 score of 1 keeps only positions where every overlapping 100-mer maps to one location in the genome. For patients LD-002 and LD-003, variants failing sequence verification during hg19 to hg38 liftover were removed.

To efficiently identify variants present in the sequencing data, we used a custom python script (python version 3.6.2), which utilizes the pysam module to efficiently extract alignments harboring variants, and extracted any read that both uniquely maps to a variant of interest and was in an aligned portion of the read (no clipping or soft-masking at the position of the variant), as well as all sequencing features for that read. Per-sample variant base quality (BQ) and mean read base quality (MRBQ) cutoffs were tuned individually for each cell-free DNA sample by single variable ROC analysis, separating reads supporting matched germline variants from reads supporting site-level sequencing errors. The thresholds maximizing germline-versus-error separation per sample were used as the BQ and MRBQ cutoffs for that sample. The per-sample coverage cutoff was set similarly, using half the median sequencing depth as a minimum-coverage floor.

Reads at the remaining variant sites were filtered per-read with the following thresholds: variant base quality at or above the per-sample BQ cutoff, mean read base quality at or above the per-sample MRBQ cutoff, MAPQ at or above 60, fragment length between 40 and 240 bp to prevent genomic DNA contamination, raw edit distance below 2, number of bases on the read with PHRED-scaled base quality below 20 at or below 5, position in read between 10 and 140, pre-denoise VAF at or below 0.2, mean reference-read edit distance at the site at or below 1, and primary alignment score (AS) minus supplementary alignment score (XS) at or above 100. All read-level metrics were computed on the individual paired-end reads. Only variants supported by reads with R1/R2 concordance were retained. Read pairs were additionally merged with FLASH2, and position in read was recomputed on the merged read. Merged reads were required to have position in read above 25 on the forward strand and below 135 on the reverse strand.

Each retained variant was further required to have VAF at or below 0.1 from the samtools mpileup, multiallelic variant proportion at or below 0.01, non-null MQSB, and depth above the per-sample coverage cutoff, defined as greater than half the median sample coverage.

For each cell-free DNA sample, the total liver detection rate was computed as the number of unique read IDs supporting the patient-specific variant set, divided by the summed per-site sequencing depth across that set.

The same patient-specific variant set was queried for three sample classes: the matched serum cell-free DNA from that patient, cross-patient serum cell-free DNA from other chronic liver disease patients, and cell-free DNA from non-liver disease controls. All three classes were processed with identical denoising filters and identical per-site depth requirements, so unmatched and control samples measure the background error rate over the same loci at the same depth.

Detection was tested per sample with a one-sided exact binomial tail on the number of supporting reads, taking the number of interrogated reads at the patient-specific sites as the trial count. The background rate was estimated specific to each patient’s VCF, since the interrogated variant set differs by patient and the control error rate is therefore patient-specific. Within each patient’s variant set, the background rate was computed as a leave-one-out pooled rate over the non-liver disease controls, so that a control sample is never tested against a background estimate to which it contributed. Matched and cross-patient samples never contribute to the background estimate.

### ppmSeq cell-free DNA SNV Detection Informed by LCM Liver Biopsies

Plasma cell-free DNA reads were processed through the FlowFeatureMapper pipeline and restricted to reads overlapping the patient-specific LCM-derived somatic mutation call set defined above. Sites falling in centromeres, simple repeats, the ENCODE blacklist regions, or the Ultima low-confidence regions were removed. Variants observed in six or more plasma samples across the cohort, tabulated from alignment-level alternate-read support rather than from called variants, were removed as recurrent plasma artifacts.

Per-read filters were applied to retain only high-confidence supporting reads: ML QUAL ≥ 2, X_SMQ ≥ 35 on both flanking regions, X_SCORE ≥ 7, edit distance ≤ 2, X_ INDEX ≥ 12, read length ≤ 220, and alignment flags restricted to 0 and 16. Only mixed reads were retained, so that every supporting read carries double-stranded support for the mutation. Each variant’s plasma alt-read count was tested with a one-sided binomial test against an in-house UG whole-genome panel (n = 91 WGS from normal tissue) for error polishing, retaining variants at p < 0.01, and variants were required to have depth above half the sample mean coverage, capped at a minimum of 20x.

Only mutations with VAF below 0.1 were used for the detection rate. For ppmSeq, the detection rate is defined as the number of unique mixed supporting reads divided by the product of the mean coverage, the number of interrogated mutations, and the mixed-read rate of the sample, since only mixed reads were used in the ppmSeq analysis.

The same patient-specific variant set was queried for the matched plasma cell-free DNA, cross-patient plasma cell-free DNA from other chronic liver disease patients, and cell-free DNA from non-liver disease controls, under identical filters and depth requirements, so that unmatched and control samples measure the background error rate over the same loci. We then applied a lasso, in which a cell-free DNA sample with only one supporting read across the entire patient-specific variant set had its detection set to zero.

Detection was tested with a one-sided exact binomial tail on the number of supporting reads against the interrogated molecule count. The background rate was estimated specific to each patient’s VCF and computed as a leave-one-out pooled rate over the non-liver disease controls, so that a control sample is never tested against a background estimate to which it contributed. Matched and cross-patient samples never contribute to the background estimate. For patient LD-001, no control retained a supporting read after the lasso, leaving a background rate of zero. We therefore bounded the background rate using the rule of three, taking 3/N as the upper limit on a rate observed as zero in N interrogated molecules, and evaluated the binomial tail at that bound.

### ppmSeq Liver Nodule Somatic Mutation Calling

Variants were identified using FlowFeatureMapper from the UG-specific GATK suite. We removed variants with a gnomAD allele frequency above 0.001, variants present in the 1000 Genomes panel of normals, and variants falling in centromeres, simple repeats, the ENCODE blacklist regions, or the Ultima low-confidence regions. Reads were required to have X_SCORE ≥ 5, X_SMQ ≥ 30 on both flanks, edit distance ≤ 4, and alignment flags of 0 or 16. Variants were then required to have at least two raw supporting reads in the nodule, along with either one MIXED read or three reads in total at ML QUAL ≥ 11. ML QUAL is a machine-learning recalibrated quality assigned to each variant-supporting read, trained to separate true variants from sequencing and library artifacts using the read’s flow signal, strand representation and alignment features.

To remove residual germline variants, we excluded any variant with a mean VAF above 0.35 across nodules, and any variant whose VAFs across nodules were not overdispersed under a beta-binomial fit (rho < 0.1), consistent with previous studies calling somatic mutations across multiple tissue biopsies from a single donor^15^.

To suppress background sequencing noise, we built an empirical error model from a Panel of Normals. At each candidate site, we computed both an alt-specific error rate (PON alt-supporting reads over PON depth at that specific mutation) and a site-level error rate (PON alt-supporting reads over PON depth at that locus, aggregated over all non-reference bases). For each variant, we tested the observed alt-read count against both error rates using a one-sided binomial test (alternative: observed greater than expected under the error rate) and required p < 0.05 against both the alt-level and the site-level model. We further required the alt-level error rate and site-level error rate to each fall below 0.01 and required the VAF to exceed the alt-level error rate by at least fivefold. In parallel, we constructed a nodule-level blacklist by tabulating, for each candidate variant, the number of distinct nodules across the full liver cohort in which it was observed; variants present in 20 or more nodules across our WGS nodule database were considered recurrent calls and removed.

### ppmSeq cell-free DNA SNV Detection Informed by Liver Nodule Biopsies

Plasma cell-free DNA from the cirrhosis patients was processed exactly as described in “ppmSeq cell-free DNA SNV Detection Informed by LCM Liver Biopsies”, with the same blacklist region exclusions, per-read filters, plasma recurrence exclusion, binomial testing and depth floor. The only difference is the input call set, which here is the nodule-derived somatic mutation call set defined in “ppmSeq Liver Nodule Somatic Mutation Calling”. Variants nominated in more than one liver donor were excluded as artifacts.

Only mutations private to a single nodule were used, to prevent infiltration of clonal hematopoiesis and germline mutations into the patient specific set of chronic liver disease mutations. The detection rate is the number of unique supporting mixed reads divided by the product of the mean coverage, the number of interrogated mutations, and the mixed-read rate of the cell-free DNA sample. We derived a total liver rate per patient, pooling all private mutations across nodules, and a per-nodule rate, mapping each mutation back to its nodule to calculate a nodule-specific shedding rate in the cell-free DNA.

The same patient-specific variant set was queried for the matched plasma cell-free DNA, cross-patient plasma cell-free DNA from other chronic liver disease patients, and cell-free DNA from non-liver disease controls. We then applied the same lasso, in which a cell-free DNA sample with only one supporting read across the entire patient-specific variant set had its detection set to zero.

### Spatial Mapping of cell-free DNA Detection onto Liver Architecture for LD-001

To ask whether cell-free DNA contribution varied across the liver, we computed detection per biopsy using only variants unique to that biopsy, so each biopsy is scored on mutations not shared with any other biopsy from the same patient. For patient LD-001, we mapped these rates onto the phylogeny built from somatic SNVs across nodules of liver Segment E and onto the matched hematoxylin and eosin-stained section.

### Biopsy-Naive Plasma ppmSeq Variant Calling and Mutational Signature Analysis

To identify somatic mutations in cell-free DNA without matched liver sequencing, variants were called *de novo* from each cell-free DNA featuremap. ML QUAL was thresholded per trinucleotide context in the orientation read by the sequencer and separately for each sample: within each context we computed a receiver operating characteristic curve using ML QUAL to separate germline from rare positions and took the threshold maximizing Youden’s J statistic. Variants were required to be supported by a mixed read with X_SMQ ≥ 35 on both flanks, X_ SCORE ≥ 7, edit distance ≤ 2, X_INDEX ≥ 12, read length ≤ 220, alignment flags of 0 or 16, and a FeatureMap Filter Status of PASS. Analogous to biopsy informed mutation calling, we removed variants in centromeres, simple repeats, the ENCODE blacklist regions, and the Ultima low-confidence regions.

Each variant was assigned its trinucleotide context from the GRCh38 reference and pyrimidine-normalized to the 96 single-base substitution classes. Per-sample 96-class counts were refit against a fixed catalog using MuSiCal^75^ (likelihood_bidirectional, threshold 0.01). The catalog comprised a clonal hematopoiesis signature as a control^47^, SBS16 for alcohol exposure, and the de novo MASLD signature previously identified in MASLD liver biopsies^15^, assembled from COSMIC v3.3.1. Exposures are reported as the fraction of each sample’s total assigned burden.

### Fragment length analysis at somatic mutation sites

To compare the size distribution of liver-derived and non-mutant cell-free DNA molecules in ppmSeq, we queried the matched plasma at the genomic positions of somatic mutations identified in matched liver sequencing. At each position we took the post-denoised mixed reads supporting the mutation, and every read carrying the reference allele at that position, requiring MAPQ 60 and excluding secondary, supplementary and duplicate alignments. ppmSeq reads are single-end, so fragment size was taken as the aligned read length. Reads longer than 220 bp were excluded, as were mutations with plasma coverage at or below 20x. Fragment sizes were pooled across all positions and all patients, and mutant and reference distributions were compared with a two-sided Wilcoxon rank-sum test.

We applied the same methodology to two mutation sets. Liver-derived mutations were those present in a single nodule, restricted to nodules with a lasso nodule-unique detection rate greater than zero in the matched cell-free DNA. Clonal hematopoiesis mutations were those present in every nodule of a patient, with no nodule restriction.

We applied the same methodology to plasma cell-free DNA from bladder cancer patient BLA-329, sequenced with ppmSeq, using mutations identified from the matched tumor as the mutation set (**Supplementary Fig. 3A**).

For the Illumina serum cell-free DNA from patients LD-001, LD-002 and LD-003, mutation-supporting reads were taken from the denoised read set defined in “Biopsy-Informed cell-free DNA SNV Detection in LCM-Profiled Livers (Illumina)”, and non-mutant reads were taken from the aligned BAM file at the same positions, restricted to primary, non-duplicate, properly paired reads with an aligned base at the site and a fragment size between 40 and 240 bp. Fragment sizes were extracted from the TLEN field, pooled across the three patients and compared with a two-sided Wilcoxon rank-sum test (**Supplementary Fig. 4**).

### Targeted Sequencing Panel Design

To leverage the opportunity to non-invasively monitor cell-free DNA bearing liver-derived clonal mutations in liver disease patients, we took an expansive approach to panel design to broaden the opportunity for chronic liver disease-associated gene detection. Our targeted panel included 17 genes as chronic liver disease-associated somatic mutation (CASM) genes (**Supplementary Table 4**). These comprised genes that were identified from previous data as being somatically mutated in diseased livers^15,16,60^ (*FOXO1, ACVR2A, CIDEB, GPAM, TNRC6B, CHD4, FASN, INSR, A1CF*). The remaining genes (*KLF15, UBE2D3, CDKN1B, SF3B3, EEF2, TNFRSF10B, HPN, ADH1B*) were included based on data from the Campbell lab nominating these genes as chronic liver disease drivers (Brzozowska et al., manuscript in preparation). The panel additionally included a set of genes to use as comparators and controls in the development of our chronic liver disease-specific detection panel. These genes included clonal hematopoiesis drivers^6,49^, (*DNMT3A, TET2, ASXL1, TP53*), hepatocellular carcinoma drivers^50^ (*CTNNB1, AXIN1, ARID2, BAP1*), inflammatory bowel disease drivers^21–23^ (*ETV6, FBXW7, HNRNPF, KRAS, IL17RA, NFKBIZ, NOS2, PIGR, RNF43, TRAF3IP2, ZC3H12A*), colorectal cancer drivers^51^ (*APC, PIK3CA, SOX9, SMAD4*) and genes that are specifically highly expressed in the liver^15,48^ (*ALB, TF, APOB, CYP2E1, FGA, FGG, FGB, CYP2C8, SERPINA1*).

### Targeted Panel cell-free DNA and Liver Nodule Preprocessing

Cell-free DNA libraries were prepared with IDT fixed unique molecular identifiers (UMI) adapters and processed using the fgbio toolkit (https://github.com/fulcrumgenomics/fgbio), following developer guidelines for duplex consensus calling. Briefly, the IDT fixed UMI was extracted using ExtractUmisFromBam, and then converted back to fastqs using SamToFastq. To prevent adapter contamination, skewer (v2.1) was used to trim adapters. Reads were then aligned to the human reference genome (hg38), and fixed IDT UMIs were corrected with CorrectUMIs (max-mismatches 1 and min-distance 2, as per fgbio best practices), and grouped by UMI using GroupReadsByUMI with parameters -m 0 -s paired -e 1. Single strand and duplex consensus sequences were created using CallMolecularConsensus and CallDuplexConsensus, then remapped to the human reference genome (hg38), integrating consensus read level tags with MergeBamAlignment. Finally, variant allele fractions were calculated using Lofreq.

#### Cell-free DNA SNV calling

### Overview and inputs

The yosemite pipeline (see Code Availability) performs SNV calling on duplex- and single-strand consensus sequence (SSCS)-aligned BAMs produced upstream by the yellowstone pipeline (see Code Availability). yosemite is implemented in Nextflow (DSL2) with per-process containerization and uses LoFreq duplex-consensus calls as the candidate variant set, which is then re-genotyped, denoised, and tested against a bootstrapped, hierarchical empirical Bayes Beta-Binomial model fit to a panel of normals.

### Library size and depth tabulation

For each sample, per-gene library size was computed separately for the duplex and SSCS BAMs using bedtools multicov against the panel target BED. Per-position sequencing depth was computed for both consensus types using samtools mpileup restricted to panel-targeted positions.

### Candidate extraction and per-read feature collection

For each candidate variant in the duplex-consensus LoFreq VCF, restricted to single-base substitutions, reads spanning the variant position were fetched in parallel per chromosome from both the duplex-consensus and SSCS realigned BAMs using pysam. Secondary, supplementary, duplicate, and improperly paired reads were discarded. For cell-free DNA samples, an additional template-length cap of 700 bp was applied to remove implausibly long fragments. For each retained read, we computed position-in-read (PIR) with CIGAR-aware soft-clip correction, base quality at the variant position (BQ), mean read base quality (MRBQ), fragment length, MAPQ, edit distance (raw and normalized to alignment length), number of low-quality positions (BQ < 20), N count, template orientation, read-pair orientation, and the duplex and single-strand consensus tags from the fgbio-style consensus calling (aD, bD, cD; aE, bE, cE; aM, bM, cM; ad, bd, cd; ae, be, ce; ac, bc; aq, bq; UQ), CIGAR string, and the MI and RX UMI tags. The minimum distance from fragment end was computed from PIR, fragment length, and read length, using the duplex (ad/ae) or single-strand (cd/ce) consensus length arrays as appropriate.

For each candidate variant, reads supporting the alternate allele were classified as concordant (alternate base observed on both mates of the read pair), discordant–non-overlapping (alternate observed on only one mate, with no overlapping ref or error call on the other), or true-discordant (alternate observed on one mate with the reference or a third-allele error on the other).

### Read position bias

For each LoFreq candidate, read position bias (RPB) was computed in parallel using pysam: for every read overlapping the variant, the normalized in-read position (query position divided by read length) was recorded separately for reads supporting the alternate and reference alleles. The two distributions were compared with the Wilcoxon rank-sum test, and RPB was reported on the PHRED scale (−10 log10 P), as in bcftools.

### UMI Family Statistics

For each variant in the sample’s duplex LoFreq VCF, the underlying UMI families were re-inspected from the pre-consensus BAM to assess consensus reliability. A duplex consensus inherits the alignment and base-quality properties of its constituent reads, and UMI families with frequent indels, soft clips, or elevated edit distance can support spurious consensus calls. Per UMI family, we computed the number of reads with indels, the number with soft-clipped bases, mean soft-clip length, mean number of low-base-quality positions, and mean edit distance stratified by read-pair orientation. These statistics were computed at two resolutions matched to the denoising stage at which they are applied: at the duplex level, pooling the two strands of a confirmed duplex pair into a single family, for use when filtering duplex-supporting reads; and at the single-strand level, treating each strand independently, for use when filtering single-strand consensus reads that lack a duplex mate.

### Duplex molecular-tag uniqueness filter

To remove residual library-preparation artefacts, the same mutation supported by multiple molecular index (MI) tags whose read 1 start positions, read 2 start positions, and fragment lengths matched within 2 bp (across both strand orientations) was flagged as a likely PCR or capture artefact, and the offending MI tags were removed from the variant-supporting set. This filter handles cases where a single template molecule is incorrectly tagged with multiple MI values.

### Duplex- and Single Strand Consensus (SSCS) -level read filters

All reads (duplex and SSCS) with insertion or deletion events in the CIGAR or with 5’-clipped CIGAR operators were removed. For each read family, base-level error rates at the variant position were parsed from the duplex error arrays (Vad, Vae, Vbd, Vbe) and single-strand error arrays (Vcd, Vce), and per-read sums of strand-specific error rates across the read were retained as features.

Variant-supporting reads were denoised in two stages. Duplex reads were filtered first. SSCS reads were filtered next, after excluding any read belonging to a denoised duplex family.

A duplex read was retained if the per-strand consensus error rate at the variant base was zero on both strands, Variant BQ ≥ 60, PIR between 18 and 125, distance from fragment end > 18 bp, edit distance ≤ 2, MAPQ ≥ 40, read family consensus error rate ≤ 0.05, and UMI-family mean low-quality-base count ≤ 5. The UMI family was also required to be free of indel-bearing reads, and reads with discordant calls between paired mates or with variants present in gnomAD or dbSNP were removed. A mutation was carried forward if more than one duplex read supported it, or if a single supporting duplex read came from a duplex family of more than two underlying reads (cD > 2). For cell-free DNA libraries, at least one supporting read per mutation was further required to come from a fragment ≤ 320 bp to prevent genomic DNA contamination.

Single-strand consensus reads were retained if BQ ≥ 30, MAPQ > 40, PIR between 18 and 125, distance from fragment end > 18 bp, edit distance ≤ 2, per-read low-quality-base count ≤ 10, no single-strand consensus disagreement at the variant base, and cE ≤ 0.05. The same UMI-family, paired-mate discordance, and germline filters applied as for duplex reads.

For each variant, we then tabulated the number of pre-denoise duplex reads, the number of duplex reads passing denoising, and the number of simplex reads passing denoising, which together form the supporting-read counts used as inputs to the downstream variant calling filters.

### Panel-level blacklist filters

Candidate mutations were intersected against three pre-computed panel-level blacklists, each of which removes mutations from the candidate set. The first is a mappability blacklist derived from a 100-bp k-mer simulation across the panel: mutations at positions where the simulation flagged the k-mer as ambiguously mappable were removed, retaining only mutations at uniquely mappable positions. Recurrent background artefacts were removed using two blacklists specific to our targeted sequencing space, the first derived from non-denoised simplex calls across a database of 301 cell-free DNA samples and the second from non-denoised duplex calls across a database of 45 liver nodule gDNA samples.

### Background allele suppression

To distinguish low-VAF somatic variation from sequencing and library-preparation noise, candidate mutations were tested against a hierarchical empirical Bayes beta-binomial model of background noise fit to a panel of normals (PoN). The panel comprised the 46 non-liver-disease control samples used throughout this study, after exclusion of samples with suspected chronic liver disease. Within each control, non-reference alleles present at VAF > 5% were masked as germline or contamination and did not contribute to the background. The 202,287 panel sites were expanded across the three non-reference alleles per site to give 606,861 candidate mutations, cached as a mutation × sample matrix of alternate counts and depths, alongside a site × sample matrix summing alternate counts across the three non-reference alleles at each site.

Background counts were computed after excluding, independently at each position, the single PoN sample with the highest VAF at that position. Because the exclusion is applied per position, different controls are dropped at different positions. This limits the contribution of individual controls carrying uncaught germline variants or contamination at a given site.

The background model combined a panel-wide prior with two levels of local pseudocounts. Two beta priors were fit by method of moments, one on per-mutation and one on per-site alternate-allele rates, aggregated across all panel positions after the per-position exclusion above. Positions with zero background rate were retained in the fit, so that the prior describes the marginal rather than the conditional-on-nonzero rate distribution; the prior concentration was left unbounded. Local pseudocounts were then added, aggregating PoN alternate and depth counts across the ±25 bp window centered on the candidate position, with the same window counts entering both the mutation-level and site-level posteriors. Site-specific pseudocounts were added from the candidate position itself, taken at the candidate alternate allele for the mutation-level test and summed across the three non-reference alleles for the site-level test. Supporting reads for each candidate were counted from pre-denoise SSCS reads passing base quality ≥ 30 and with a PIR between 10 and 133 and tested against the resulting beta-binomial posterior with SSCS depth as the trial count. This yielded a mutation-level P value at the candidate alternate allele and a site-level P value across all alternate alleles at the position, together with the ratio of sample VAF to PoN VAF at each level.

### Bootstrap panel composition and final SNV calls

Prior fitting and beta-binomial testing were bootstrapped over PoN subsamples to prevent any single panel composition from driving a call. When the sample being called was itself one of the 46 controls, it was removed from the panel before bootstrapping, so that no iteration could draw the sample into its own background; the eligible pool was therefore 45 for control samples and 46 for all other samples. In each of 10,000 iterations, 16 PoN samples were drawn without replacement from the eligible pool, both priors were refit, local and site-specific pseudocounts were recomputed, and mutation- and site-level P values and VAF ratios were recalculated. Iteration *i* was seeded as (master seed + *i*), making calls bit-identical across runs on identical inputs.

Variant candidates were required to be supported by at least one denoised duplex read. A candidate passed an iteration under any of four criteria: (i) more than one denoised duplex read, with mutation- and site-level P < 0.05; (ii) more than one pre-denoise duplex read, with mutation- and site-level P < 0.05 and both VAF ratios > 2; (iii) at least one denoised SSCS read, with mutation- and site-level P < 0.05 and both VAF ratios > 2; or (iv) at least two supporting SSCS reads, with mutation- and site-level P < 0.05 and a site-level VAF ratio > 3. Candidates passing in at least 95% of iterations were retained.

Retained candidates were further required to have either at most 10 PoN samples non-zero at the position, averaged across iterations, or a duplex VAF at least tenfold above the highest PoN VAF at that position. Candidates supported by a single pre-denoise duplex read with no denoised SSCS reads were additionally required to have duplex VAF > 0.001. Candidates with PHRED-scaled position bias ≥ 60 were removed. A VAF ceiling of 0.1 was applied to cell-free DNA samples and 0.25 to gDNA samples, except at the clonal hematopoiesis driver genes *DNMT3A, TET2, TP53* and *ASXL1*.

### Cell-free DNA indel calling Overview and inputs

The yosemite-indels pipeline (see Code Availability) performs short-insertion and -deletion (indel) calling on the same duplex- and SSCS-aligned BAMs used by yosemite.

It is implemented in Nextflow (DSL2) and, for each sample × consensus mode, (i) filters the realigned BAM to reads carrying CIGAR insertion or deletion operations, (ii) extracts per-read indel features into an ANNOVAR-compatible avinput table, (iii) annotates with ANNOVAR table_annovar.pl (refGene, dbnsfp42c, cosmic70, avsnp150, exac03, clinvar_20220320, gnomad40_exome, ALL.sites.2015_08; hg38), (iv) reheaders the multianno output, and (v) intersects with the panel BED. Per-sample indel calling is then performed downstream.

### Indel-bearing read extraction and per-read feature collection

For each sample × consensus mode, reads carrying at least one insertion (CIGAR I) or deletion (CIGAR D) operator were extracted from the realigned BAM using pysam, sorted, and indexed. For every indel event in every retained read, we recorded: chromosome, reference start and end coordinates, reference and alternate alleles (with ANNOVAR-style “-” notation for inserted or deleted bases), indel length, MAPQ, proper-pair, secondary, supplementary and duplicate flags, alignment-score tags (AS, XN, Xn, nM, NM), fragment length, distance from read edge, surrounding base quality (mean BQ across the 10 bases flanking the indel-anchor), homopolymer length at the event position (computed by scanning the reference for the maximal run of the indel-anchor base over the ±10-bp window), variant base quality (mean BQ across the aligned region), original base quality string (OQ tag), relative read position, number of low-quality positions (BQ < 20), number of soft-clipped bases, read 1/2 identity, cross-exon flag, GC content and complexity of the 20-bp context window, indel type (insertion/deletion), fragment-orientation-corrected minimum distance from fragment end (computed with PIR both with and without soft clips), the trinucleotide context (96-class encoding adapted for indels), and the full set of duplex and single-strand consensus tags (aD, bD, cD; aE, bE, cE; aM, bM, cM; ad, bd, cd; ae, be, ce; UQ; MD, MI, RX). Reads were restricted to standard properly-paired forward/reverse flags (99, 147, 83, 163).

### Annotation and panel intersection

The per-read indel table was emitted in ANNOVAR avinput format and annotated against the hg38 build of refGene, dbnsfp42c, cosmic70, avsnp150, exac03, clinvar_ 20220320, gnomad40_exome, and ALL.sites.2015_08. The multianno output, which packs avinput per-read features into the Otherinfo column, was reheaded by merging the column-header line with the first data row (the inline subheader carrying the avinput field names) to produce a single, machine-readable header. The resulting table was intersected with the panel BED using bedtools intersect (-wa -u), preserving panel positions and dropping off-panel events.

### Duplex Denoising for Indel Candidates

For duplex-supporting reads, we required, for both cell-free DNA and gDNA samples: orientation-corrected minimum distance from fragment end (without soft clips) > 18 bp; variant base quality ≥ 60; MAPQ ≥ 60; SSCS error rate at the variant position (cE) ≤ 0.05; PIR with soft clips strictly between 18 and 125; duplex strand-support counts aD ≥ 1 and bD ≥ 1; number of low-quality positions in the read ≤ 5; and alignment score AS > 110. For cell-free DNA samples only, an additional fragment-length cap of 320 bp was applied. Each retained read was then classified by joint parsing of its CIGAR string and MD tag, and only reads carrying exactly one indel event with no substitution mismatches were retained.

### Global indel blacklists and final denoising filters

Candidate indels were intersected against a database of 301 cell-free DNA samples. For each candidate indel, we recorded both the number of cell-free DNA samples carrying the exact indel allele and the number carrying any indel at that position, allowing recurrent technical artefacts to be flagged either at the allele level or at the locus level. Final calls were restricted to indels with SSCS VAF < 10%, duplex VAF > 0.05%, and at least one supporting duplex UMI family. To suppress recurrent technical artefacts, indels were required to recur in fewer than 20 cell-free DNA samples both as the same allele and as any indel at the position. Indels longer than 20 bp were excluded, as short-read mapping becomes unreliable at this length, and indels falling within a homopolymer run of two or more bases were excluded to avoid polymerase-slippage artefacts.

### Pathogenic Variant Selection and Gene Set Fraction

For each sample, the final SNV calls from yosemite and the final indel calls from yosemite-indels were combined into a single variant set, from which pathogenic variants were selected. In CASM, hepatocellular carcinoma, IBD, and colorectal cancer driver genes, pathogenic variants were defined as SNVs with CADD PHRED > 25 or AlphaMissense observed score > 0.564, plus all indels supported by a denoised duplex read. *FASN* SNVs were exempt from this pathogenicity threshold, since prior data show that *FASN* driver SNVs do not enrich at high pathogenicity scores and applying the threshold would filter out genuine *FASN* driver signal^15^.

For each sample, the pathogenic-fragment fraction in a given gene group was computed as the number of SSCS supporting reads with BQ > 30 and PIR between 10 and 133 across pathogenic variants in that group, divided by the total reads in panel target regions for the genes in that group. Normalization by target-region depth was necessary because raw supporting-read counts are not comparable across samples given the variation in sequencing depth across the cohort. The denominator was obtained from bedtools multicov over the panel target BED, counting alignments per region from the single strand consensus BAM, and summed across the genes in each group. For CASM genes, the *FOXO1p*.*S22W* locus depth was added back into the denominator, so that the canonical hotspot contributes to the read-count normalization despite the restriction of *FOXO1* to that single site.

### Convergence of cell-free DNA Detections with Nodule-Level Clonal Convergence

A pathogenic mutation was counted as independent if it was observed in exactly one nodule of that patient. Mutations observed in two or more nodules of the same patient were excluded as inherited from a shared ancestor, with the exception of the *FOXO1* p.S22W hotspot. *FOXO1* was restricted to p.S22W and to single-base substitutions; indels were included for all other genes.

Each of the 17 CASM genes in each patient was assigned a convergence weight equal to the fraction of that patient’s nodules carrying an independently acquired pathogenic mutation in that gene. The observed statistic is the sum of these weights over the CASM genes detected in that patient’s cell-free DNA, summed across all four patients.

In each of 10,000 permutations at seed 43, every patient was assigned the same number of genes as was detected in their cell-free DNA, drawn at random without replacement from the 17-gene CASM panel, and the weights of the drawn genes were summed. The permutation breaks only the pairing between the genes detected in cell-free DNA and their nodule weights. The one-sided P value was computed as (1 + the number of permutations at or above the observed value) / (1 + 10,000).

### Pathogenicity enrichment scores

For each gene, we tabulate the total number of somatic variants identified and assign each variant an AlphaMissense score^67^ and a CADD Raw Score^68^. The null set for that gene is every panel-interrogated site in that gene carrying both scores, weighted by the background error rate at that site, so that positions more likely to generate a call are drawn more often. From this null we draw the same number of mutations as were observed in that gene, with replacement, and take the mean AlphaMissense and mean CADD Raw scores of the drawn set. This is repeated 10,000 times per gene, and the P value is the fraction of draws in which the null mean exceeds the observed mean. The AlphaMissense and CADD P values are combined with Fisher’s method for combining P values and corrected across genes with the Benjamini-Hochberg procedure.

### Gene-Set Pathogenic Enrichment Ratio

Panel genes were grouped into classes by disease context: CASM genes, inflammatory bowel disease drivers, clonal hematopoiesis drivers, liver lineage-defining genes, hepatocellular carcinoma drivers, colorectal cancer drivers, and non-disease control genes. A variant was called pathogenic if its AlphaMissense score exceeded 0.564 or its CADD PHRED score exceeded 25. The same rule was applied to the observed calls and to every interrogated site in the background table.

For each gene, the expected pathogenic rate was the fraction of that gene’s interrogated sites that are pathogenic, with each site weighted by its background error rate. The expected pathogenic count for a gene was this rate multiplied by the number of calls observed in that gene. Observed and expected pathogenic counts were summed across the genes in each class, and the enrichment ratio is their quotient.

Uncertainty was assessed by bootstrap. In each replicate, patients were resampled with replacement and the per-gene call and pathogenic-call counts recomputed, and the background sites within each gene were resampled with replacement and the per-gene expected rate refit. Intervals are 95% bootstrap percentile intervals of the ratio. The P value is twice the smaller of the proportions of replicates with a ratio at or below one and at or above one, corrected across classes with the Benjamini-Hochberg procedure.

### Patient-specific Longitudinal Variant Calling (Targeted Panel)

After variant calling was performed on two samples with detection of CASM gene variants in the pre-transplant cell-free DNA, patient-specific longitudinal variant calling was performed on all variants genotyped in the pre-transplant cell-free DNA. For each post-transplant patient, a bespoke pysam script was deployed to obtain all reads with base quality greater than 30 and position in read greater than 10 bp away from the ends of the read. Variants were required to have a simplex VAF to allele-specific error rate ratio of greater than 2 to be successfully genotyped in the post-transplant cell-free DNA.

## Supplementary Figures

**Supplementary Figure 1:**
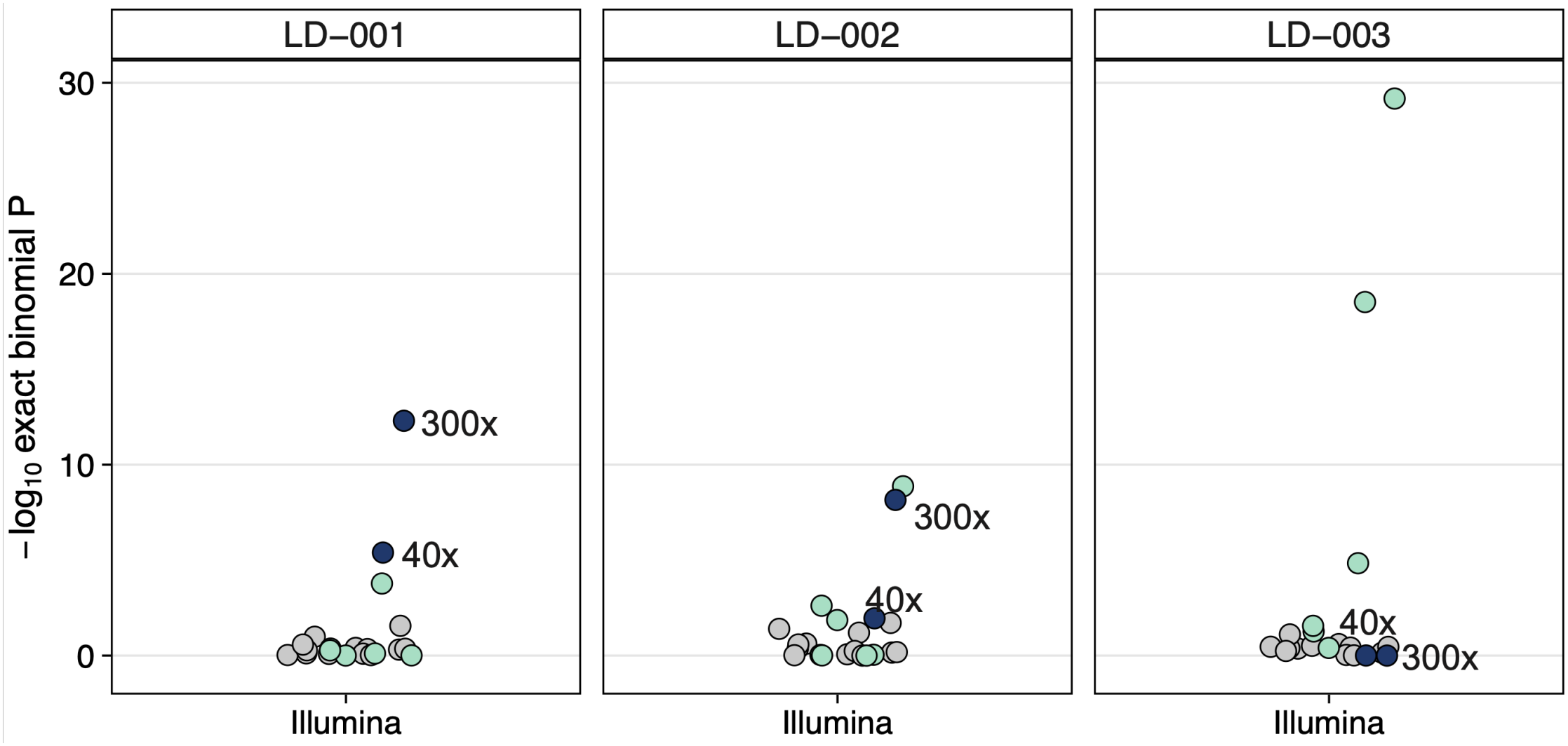
Detection of liver-derived variants using Illumina whole-genome sequencing of cell-free DNA. Binomial test P value for detected patient-specific SNVs identified from matched laser capture microdissection liver biopsies in cell-free DNA whole-genome sequences (matched serum cell-free DNA) compared to cross-patient SNV liver profiles (serum samples from n = 2 cross-patients) and cell-free DNA from non-liver disease controls (n = 13) in three patients with MASLD. Cell-free DNA was sequenced using Illumina (300x or 40x). Data are the same as those presented in Fig. 1C, but here we show 40x and 300x data for Illumina only. The bottom and top ends of the boxes represent the 25th and 75th percentiles of the data, respectively, and the horizontal lines represent the median. The whiskers represent at most 1.5 times the interquartile range.

**Supplementary Figure 2:**
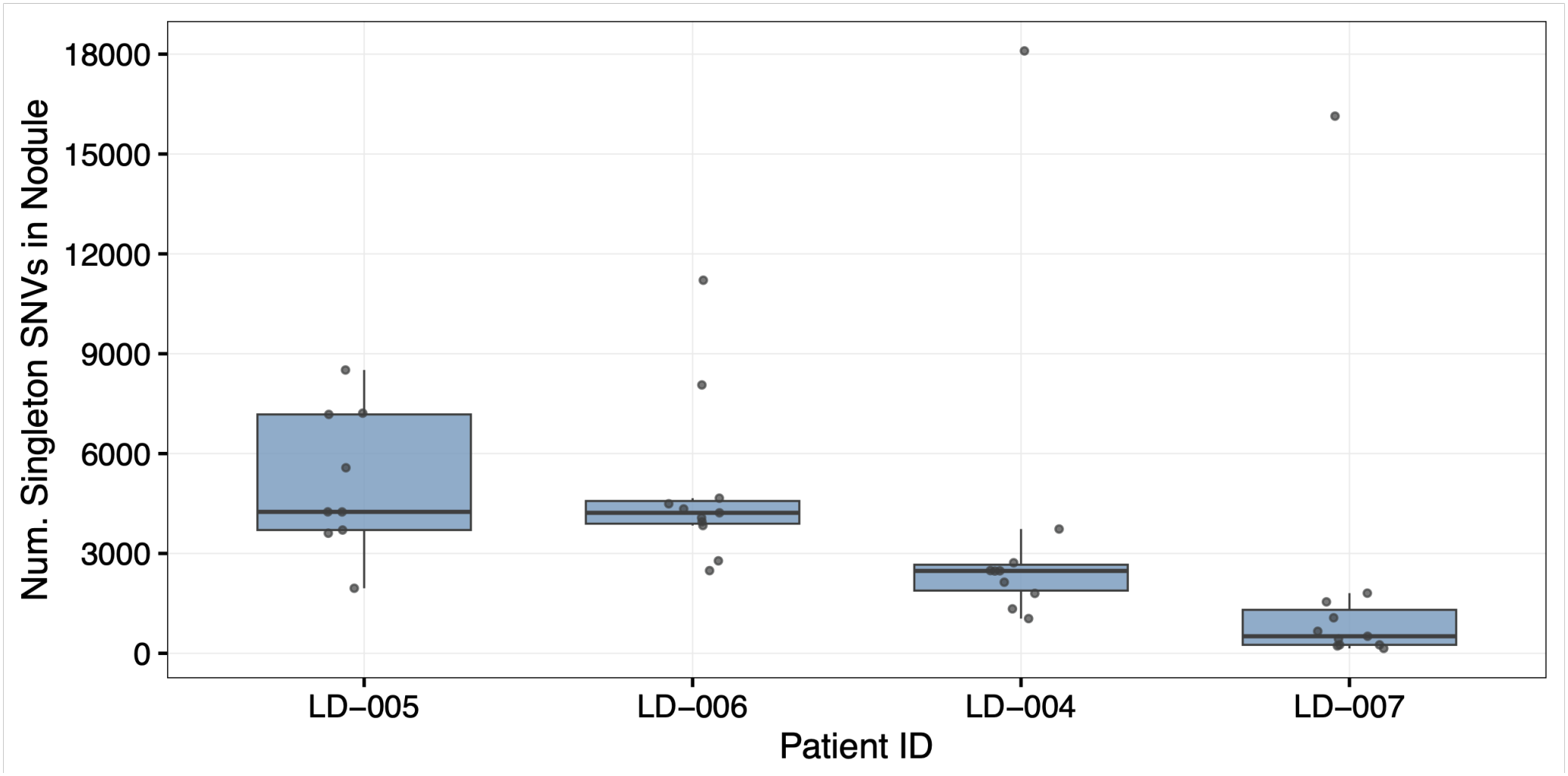
Number of somatic SNVs identified from genomic DNA sequencing of liver nodules. The number of SNVs obtained from whole-genome sequencing (ppmSeq) of genomic DNA for each liver nodule biopsy from four donors with chronic liver disease (Supplementary Table 1). Each dot represents a liver nodule biopsy (LD-005 n = 9 biopsies; LD-006 n = 11 biopsies; LD-004 n = 10 biopsies; LD-007 n = 11 biopsies). The bottom and top ends of the boxes represent the 25th and 75th percentiles of the data, respectively, and the horizontal lines represent the median. The whiskers represent at most 1.5 times the interquartile range.

**Supplementary Figure 3:**
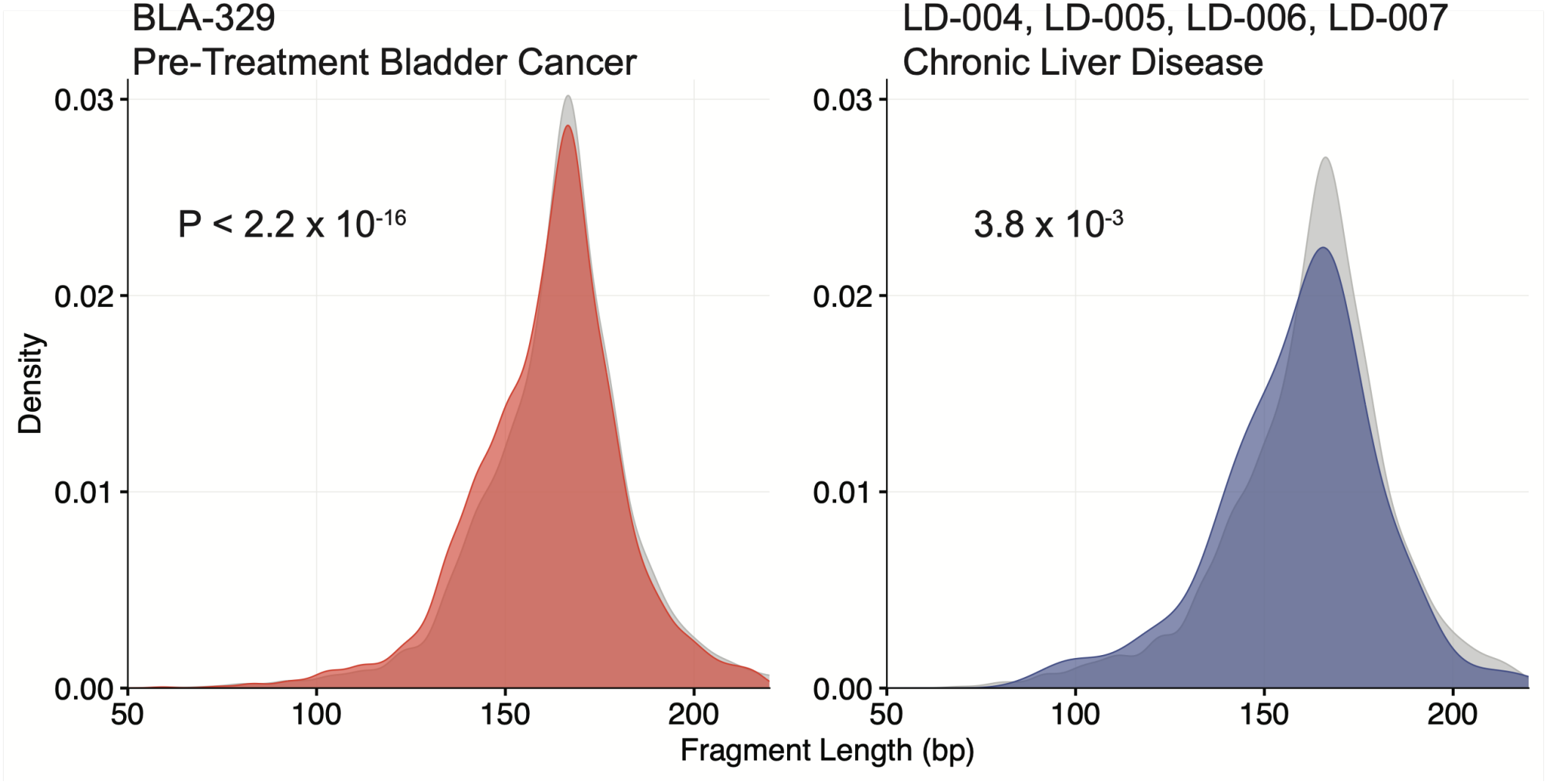
Fragment sizes of non-cancerous liver cell-free DNA mirror short fragments of circulating tumor DNA. **A**, Fragment length distribution analysis of cell-free DNA from a pre-treatment bladder cancer patient BLA-329. Fragments carrying tumor-confirmed variants identified from matched bladder tumor biopsy (ctDNA Fragment, red) compared to wild-type cell-free DNA (WT cfDNA, grey) without tumor-confirmed fragments. bp, basepairs. Two-sided Wilcoxon test. **B**, Fragment length distribution analysis of cell-free DNA for fragments carrying liver-confirmed variants identified from matched liver biopsy (liver nodule adjudicated fragments, blue) compared to wild-type cell-free DNA (WT cfDNA, grey) without liver-confirmed fragments for four patients with chronic liver disease. bp, basepairs. Two-sided Wilcoxon test. This panel is the same as the left panel in Figure 1E, shown here for comparison.

**Supplementary Figure 4.**
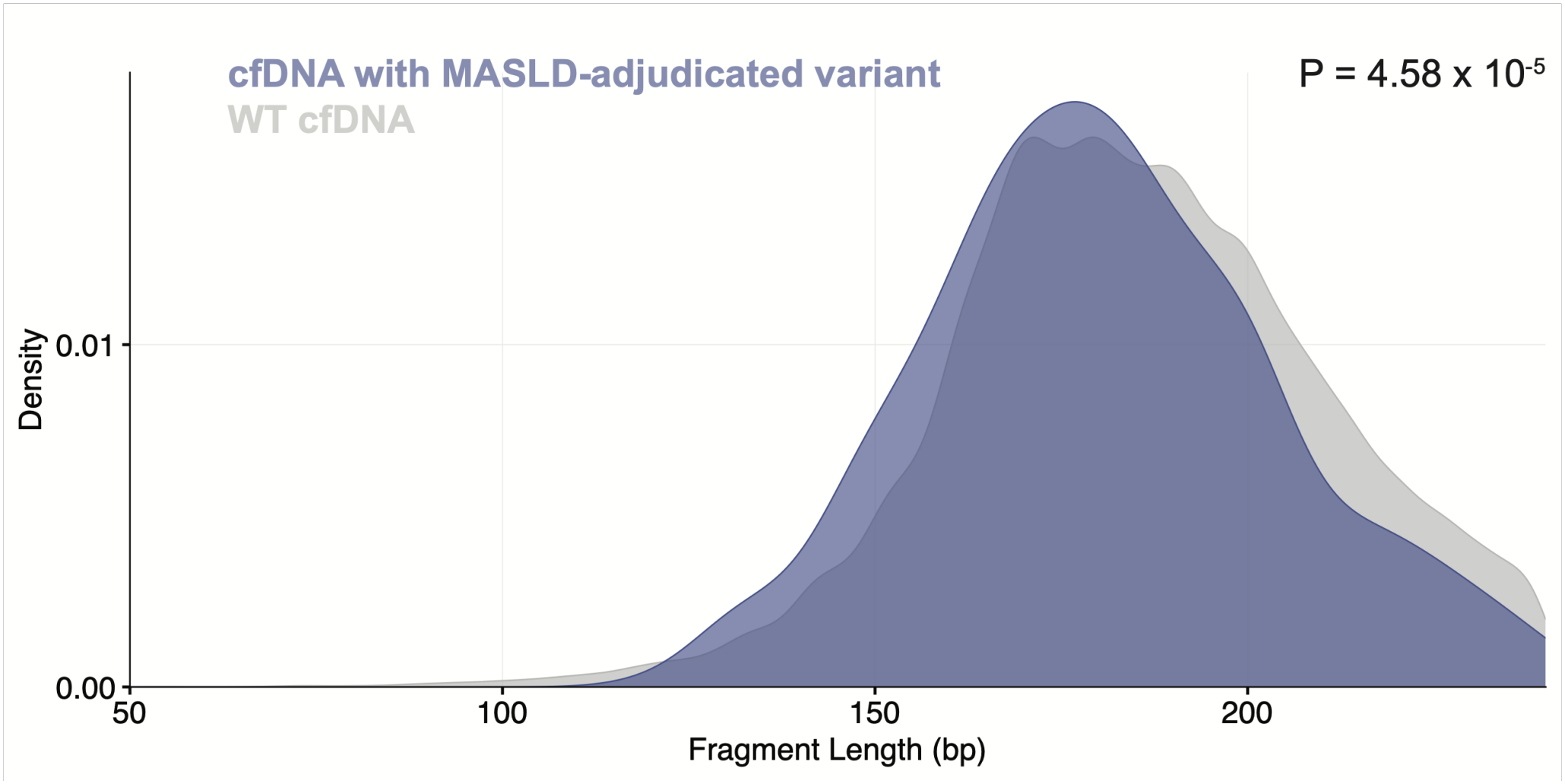
Fragment length analysis of cell-free DNA sequenced with Illumina. Fragment length distribution analysis of cell-free DNA from three MASLD patients (LD-001, LD-002 and LD-003) for fragments carrying liver-confirmed variants identified from matched liver biopsy (cfDNA with MASLD-adjudicated variant, purple) compared to wild-type cell-free DNA without liver-confirmed fragments (WT cfDNA fragment, grey), sequenced with Illumina. bp, basepairs. Two-sided Wilcoxon test.

**Supplementary Figure 5:**
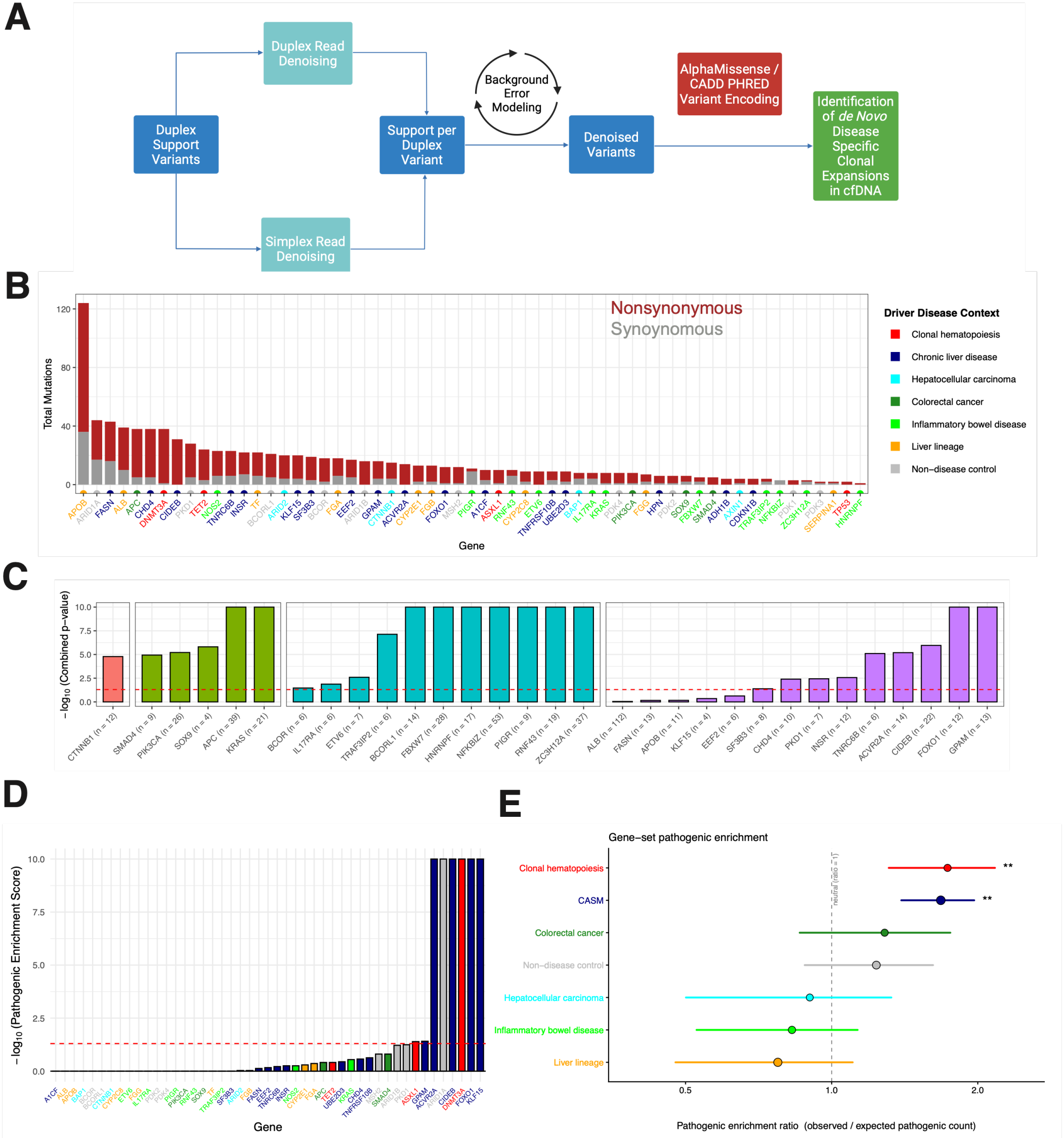
Biopsy-naive (cell-free DNA only) framework for non-invasive genotyping of chronic liver disease, cancer hotspot benchmarking analysis, and pathogenic enrichment score. **A**, Schematic of biopsy agnostic framework for variant calling. **B**, Number of total mutations for all targeted genes identified in liver nodules from four patients in Figure 2. Red indicates nonsynonymous mutations, grey indicates synonymous mutations. Genes are color coded based on their assigned disease class (blue for chronic liver-associated somatic mutation (CASM) genes, cyan for HCC drivers, orange for highly expressed liver lineage-defining genes, purple for control genes, and red for canonical clonal hematopoiesis genes). **C**, Pathogenicity enrichment scores for genes with variants previously reported by PCAWG (for HCC and CRC genes), Kakiuchi et al.^23^ (for IBD genes) and Ng et al.^15^ (for chronic liver disease genes). The number of variants in the dataset for each gene is indicated. For each gene, the total number of somatic variants observed are tabulated and each variant is assigned an AlphaMissense score^67^ and a CADD Raw Score^68^. The same number of mutations is randomly sampled across the gene, reflecting every possible mutation, to generate a null distribution. The mean AlphaMissense or CADD scores of the randomly sampled set are calculated and compared to the mean AlphaMissense/CADD scores for the detected variant set of mutations. The random sampling and mean comparisons with detected variants are repeated 10,000 times per gene. The *P* value reflects how many times out of 10,000 the randomly sampled set has a mean pathogenicity score greater than or equal to that of the observed variants. AlphaMissense and CADD scores are calculated separately, and then the two values are combined with Fisher’s method for combining p-values. The red dashed line represents the significance threshold of 0.05. **D**, Pathogenicity enrichment scores for driver genes associated with specific disease contexts calculated from somatic variants detected in liver nodule targeted panel sequencing from four chronic liver disease patients for genes with >3 variants identified. For each gene, the total number of somatic variants observed are tabulated and each variant is assigned an AlphaMissense score^67^ and a CADD Raw Score^68^. The null set for each gene is every panel-interrogated site in that gene carrying both scores, with each site weighted by the background error rate at that site, so that positions more likely to generate a call are drawn more often. The same number of mutations as observed in that gene is drawn from this null with replacement, and the mean AlphaMissense and mean CADD Raw scores of the drawn set are calculated. The sampling is repeated 10,000 times per gene. The *P* value is the fraction of draws in which the null mean exceeds the observed mean. AlphaMissense and CADD *P* values are calculated separately and then combined with Fisher’s method for combining *P* values. *P* values were corrected across genes with the Benjamini-Hochberg method. The red dashed line represents the significance threshold of 0.05. **E**, Enrichment ratio of observed versus expected pathogenic variants detected from liver gDNA as in (D) across gene classes. Points are the observed-to-expected ratio of pathogenic calls, sized by the number of mutations in that class; bars are 95% bootstrap percentile confidence intervals. Expected counts were computed per gene as the error-rate-weighted pathogenic fraction of that gene’s background sites, multiplied by the number of calls in that gene and summed within each class. P values were obtained from 10,000 two-sided percentile bootstrap replicates in which patients were resampled with replacement and every per-gene rate was refit within each replicate, then corrected across gene classes by the Benjamini-Hochberg procedure. **, q < 0.01.

**Supplementary Figure 6:**
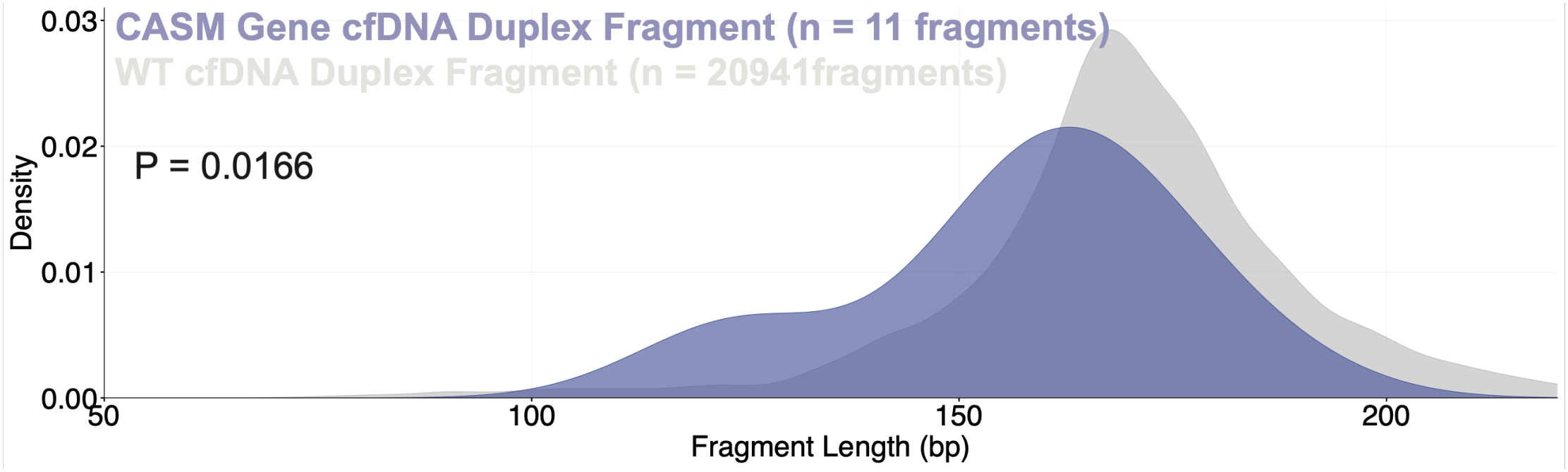
Fragment length of cell-free DNA sequenced with the targeted panel. Fragment length analysis of cell-free DNA for fragments carrying CASM gene variants (purple) compared to wild-type cell-free DNA (WT cfDNA, grey) without CASM gene variant fragments for four patients from Fig. 2B. One-sided Wilcoxon test.

**Supplementary Fig 7:**
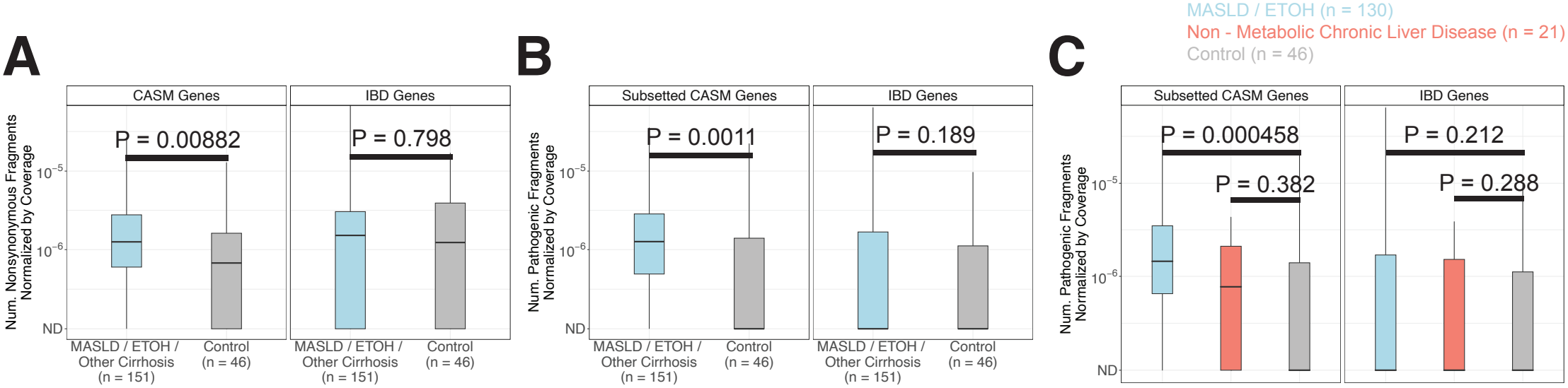
Detection of CASM gene variants in the cell-free DNA in chronic liver disease patients versus controls. **A**, Detection of nonsynonymous SNVs and indels in CASM genes (left) or IBD driver genes (right) in cell-free DNA using the targeted sequencing panel for non-chronic liver disease controls (n = 46 plasma samples) versus MASLD / alcohol / other cirrhosis (ETOH) patients (n = 151 plasma samples). Genes were assigned to one specific group (CASM or IBD). For each group, the total number of fragments with base quality >30 supporting a denoised variant is divided by the total number of duplex fragments sequenced across genes belonging to the respective group. The bottom and top ends of the boxes represent the 25th and 75th percentiles of the data, respectively, and the horizontal lines represent the median. The whiskers represent at most 1.5 times the interquartile range. Two-sided Wilcoxon test. ns, not significant. **B**, Detection of pathogenic SNVs and indels in a subset of CASM genes (*FOXO1 S22W hotspot, CIDEB, GPAM, TNRC6B, ACVR2A, FASN, CHD4, INSR*, and *A1CF*) (left) or IBD driver genes (right) in cell-free DNA using the targeted sequencing panel for non-chronic liver disease controls (n = 46 plasma samples) versus chronic liver disease (ETOH) patients (n = 151 plasma samples). This subset of CASM was identified from previous WGS/WES of chronic liver disease LCMs^15,60^ (Brzozowska et al., manuscript in preparation). For each group, the total number of fragments with base quality >30 supporting a denoised variant is divided by the total number of duplex fragments sequenced across genes belonging to the respective group. The bottom and top ends of the boxes represent the 25th and 75th percentiles of the data, respectively, and the horizontal lines represent the median. The whiskers represent at most 1.5 times the interquartile range. Two-sided Wilcoxon test. ns, not significant. **C**, Detection of pathogenic SNVs and indels in a subset of CASM genes (*FOXO1 S22W hotspot, CIDEB, GPAM, TNRC6B, ACVR2A, FASN, CHD4, INSR, and A1CF*) (left) or IBD driver genes (right) in cell-free DNA using the targeted sequencing panel for non-chronic liver disease controls (n = 46 plasma samples) versus MASLD / alcohol cirrhosis patients (n = 130 plasma samples) and other chronic liver disease conditions (n = 21). This subset of CASM was identified from previous WGS/WES of chronic liver disease LCMs^15,60^. For each group, the total number of fragments with base quality >30 supporting a denoised variant is divided by the total number of duplex fragments sequenced across genes belonging to the respective group. The bottom and top ends of the boxes represent the 25th and 75th percentiles of the data, respectively, and the horizontal lines represent the median. The whiskers represent at most 1.5 times the interquartile range. Two-sided Wilcoxon test. ns, not significant.

**Supplementary Fig 8:**
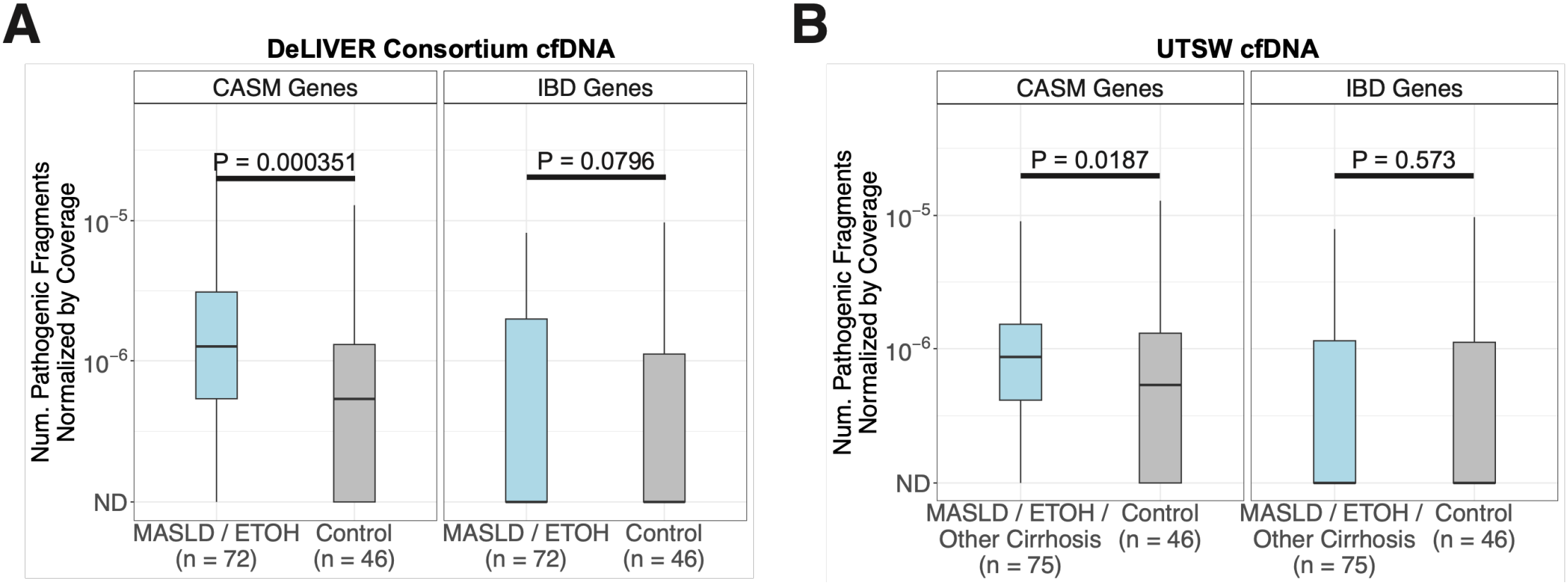
Detection of CASM gene variants in cell-free DNA stratified by sequencing center. **A**, Detection of pathogenic SNVs in CASM genes (left) or IBD driver genes (right) in cell-free DNA from the UK DeLIVER sub-cohort versus non-liver disease controls, using the targeted sequencing panel. Genes were assigned to one specific group (CASM or IBD). For each group, the total number of fragments with base quality >30 supporting a denoised variant is divided by the total number of duplex fragments sequenced across genes belonging to the respective group. The bottom and top ends of the boxes represent the 25th and 75th percentiles of the data, respectively, and the horizontal lines represent the median. The whiskers represent at most 1.5 times the interquartile range. Two-sided Wilcoxon test. **B**, Detection of pathogenic SNVs in CASM genes (left) or IBD driver genes (right) in cell-free DNA from the UTSW cirrhosis sub-cohort versus non-liver disease controls, using the targeted sequencing panel. Genes were assigned to one specific group (CASM or IBD). For each group, the total number of fragments with base quality >30 supporting a denoised variant is divided by the total number of duplex fragments sequenced across genes belonging to the respective group. The bottom and top ends of the boxes represent the 25th and 75th percentiles of the data, respectively, and the horizontal lines represent the median. The whiskers represent at most 1.5 times the interquartile range. Two-sided Wilcoxon test.

**Supplementary Fig 9:**
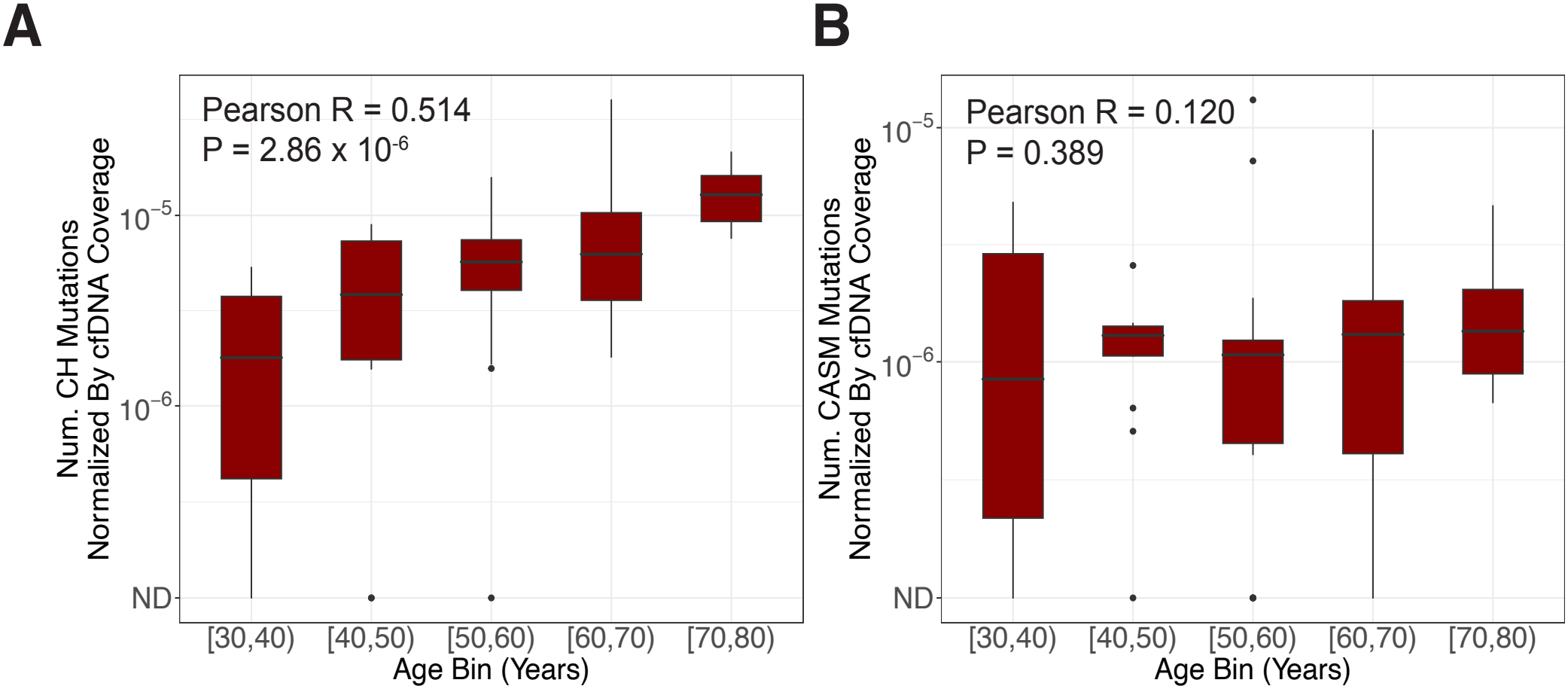
Detection of variants in clonal hematopoiesis driver genes and CASM genes from cell-free DNA across age. **A**, Number of pathogenic variants in clonal hematopoiesis (CH) panel driver genes normalized by library size from the cell-free DNA across patient age. The bottom and top ends of the boxes represent the 25th and 75th percentiles of the data, respectively, and the horizontal lines represent the median. The whiskers represent at most 1.5 times the interquartile range. Pearson’s correlation, two-sided t-test. The panel is the same as in Figure 3B, shown here for comparison. N = 74 patients with available age information were included in this analysis. **B**, Number of pathogenic variants in CASM genes normalized by library size from the cell-free DNA across patient age. The bottom and top ends of the boxes represent the 25th and 75th percentiles of the data, respectively, and the horizontal lines represent the median. The whiskers represent at most 1.5 times the interquartile range. Pearson’s correlation, two-sided t-test. N = 74 patients with available age information were included in this analysis.

**Supplementary Figure 10:**
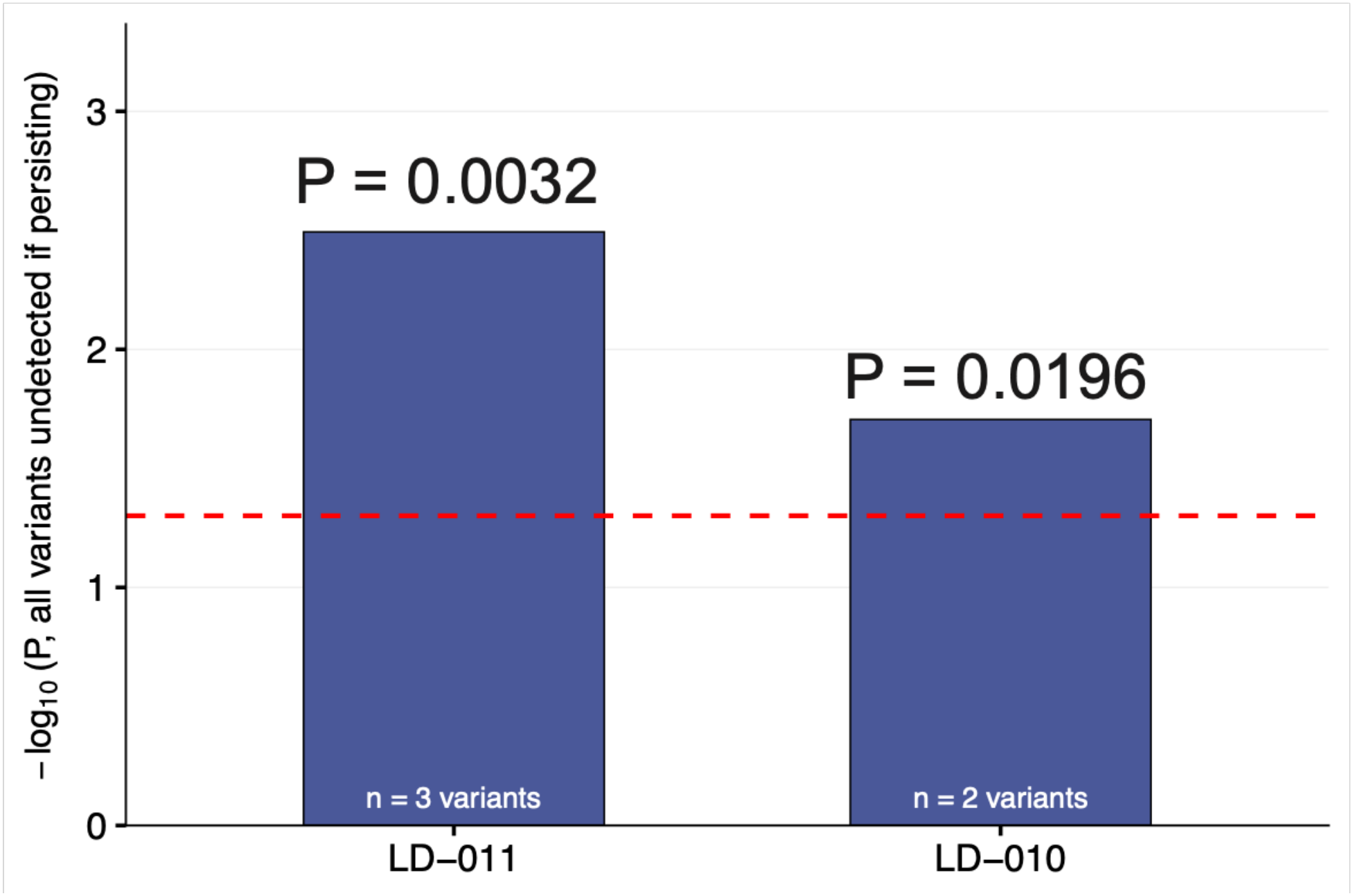
Probability of seeing zero reads for longitudinal analysis post-transplant. Power analysis showing the probability that all pre-transplant CASM drivers in a given patient would be missed post-transplant by chance alone if the clones had persisted. For each variant, the binomial probability of redetection was computed from its pre-transplant VAF and the post-transplant duplex depth, and the per-patient probability of joint non-detection was taken as the product of the per-variant non-detection probabilities across all variants in that patient.

**Supplementary Figure 11:**
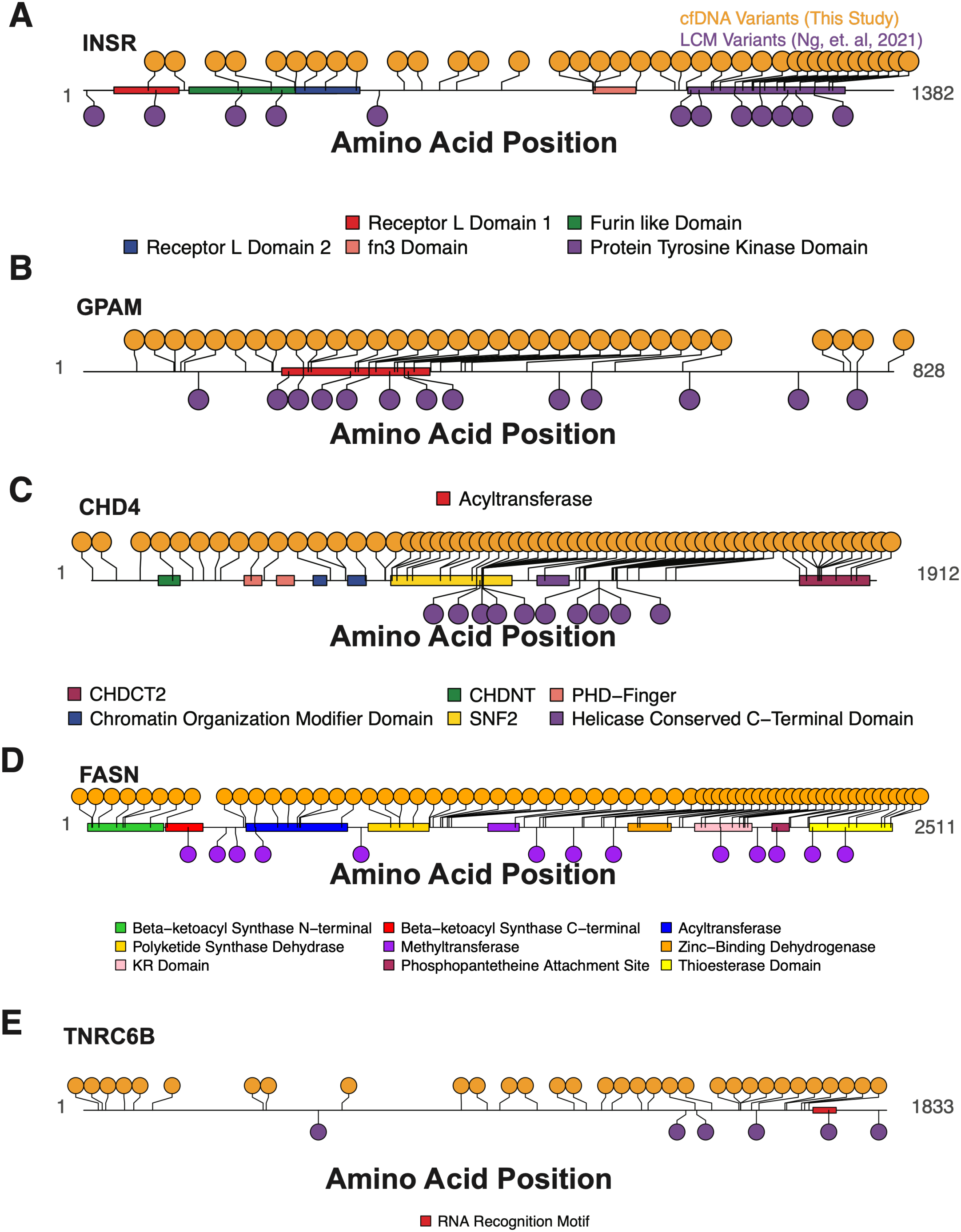
Illustration of protein domains for CASM genes and detected variants from cell-free DNA compared to previously identified variants from liver biopsies. **A**, Annotated INSR protein plot displaying cell-free DNA variants detected in the targeted panel sequencing expansion cohort (n = 151 patients) from the current study (yellow, top) and liver variants identified from whole-genome sequencing of laser capture microdissections from Ng et al.^15^ (purple, bottom). INSR domains are indicated. FN3, fibronectin type 3 domain. **B**, Same as (A) for GPAM. **C**, Same as (A) for CHD4. CHDNT, CHD4 N-terminal domain; PHD-Finger, plant-homeodomain finger; Chromo, chromodomain; SNF2, SNF2-like ATPase motor domain; CHDCT2, CHD C-terminal 2 domain **D**, Same as (A) for FASN. KR, ketoacyl reductase. **E**, Same as (A) for TNRC6B.

**Supplementary Figure 12:**
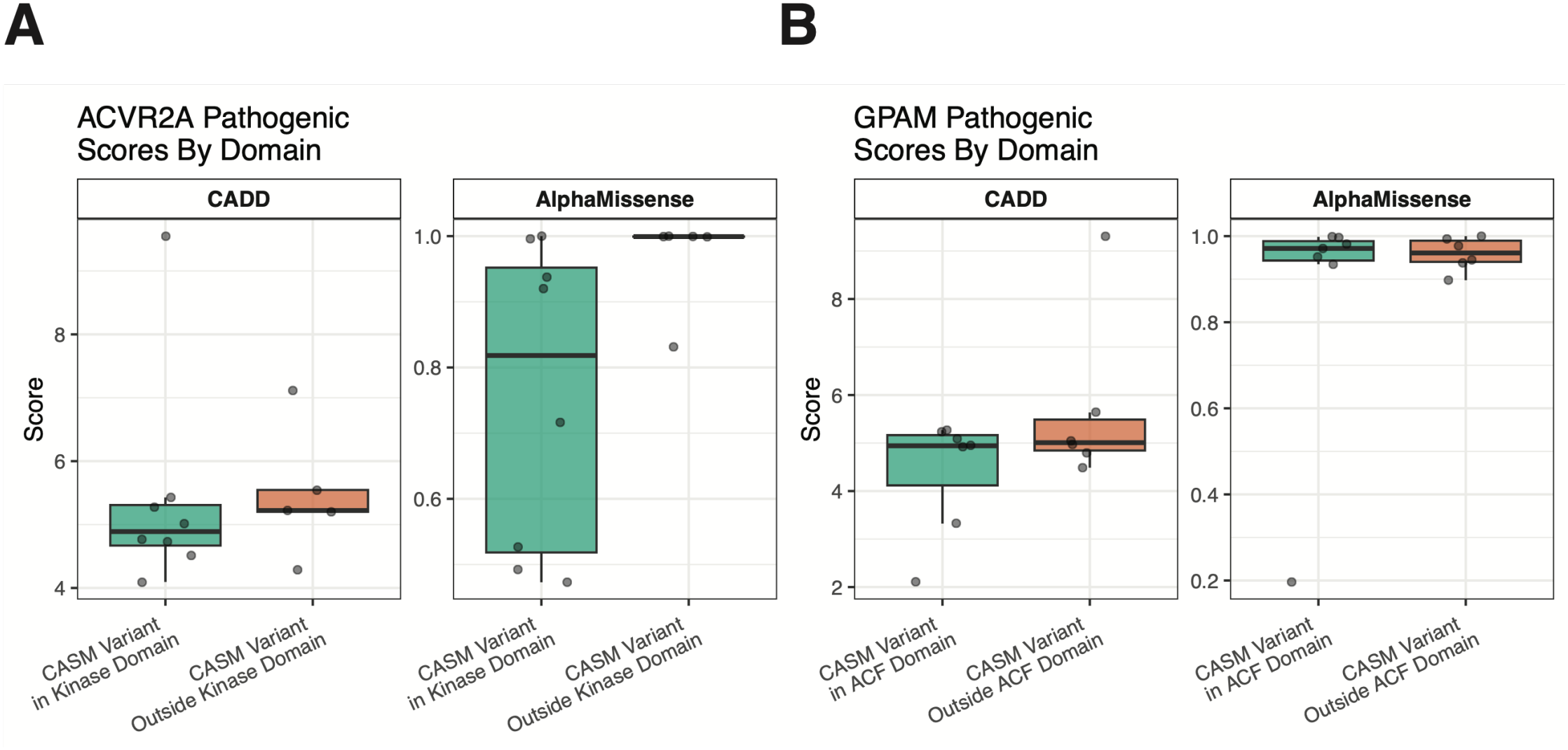
Pathogenicity scores of variants from MASLD biopsies stratified by domains. **A**, AlphaMissense and CADD scores of *ACVR2A* somatic variants in MASLD Biopsies from Ng et al. (*Nature* 2021)^15^, stratified by location in Kinase domain. The bottom and top ends of the boxes represent the 25th and 75th percentiles of the data, respectively, and the horizontal lines represent the median. The whiskers represent at most 1.5 times the interquartile range. **B**, AlphaMissense and CADD scores of *GPAM* somatic variants in MASLD Biopsies from Ng et al. (*Nature* 2021)^15^, stratified by location in Acyltransferase (ACF) domain. The bottom and top ends of the boxes represent the 25th and 75th percentiles of the data, respectively, and the horizontal lines represent the median. The whiskers represent at most 1.5 times the interquartile range.

